# UniPath: A uniform approach for pathway and gene-set based analysis of heterogeneity in single-cell epigenome and transcriptome profiles

**DOI:** 10.1101/864389

**Authors:** Smriti Chawla, Sudhagar Samydurai, Say Li Kong, Zhenxun Wang, Wai Leong Tam, Debarka Sengupta, Vibhor Kumar

## Abstract

Here, we introduce UniPath, for representing single-cells using pathway and gene-set enrichment scores by a transformation of their open-chromatin or gene-expression profiles. Besides being robust to variability in dropout, UniPath provides consistency and scalability in estimating gene-set enrichment scores for every cell. UniPath’s approach of predicting temporal-order of single-cells using their gene-set activity score enables suppression of known covariates. UniPath based analysis of mouse cell atlas yielded surprising, albeit biologically-meaningful co-clustering of cell-types from distant organs and helped in annotating many unlabeled cells. By enabling unconventional analysis, UniPath also proves to be useful in inferring context-specific regulation in cancer cells.

## Introduction

Single-cell RNA sequencing (scRNA-seq) and single-cell open-chromatin profiling help us to decipher cellular heterogeneity of activity of coding and non-coding genomic elements[1, 2]. The heterogeneity in the activity of genomic sites among single-cells is regularly used to estimate cellular composition in complex tissue, spotting rare cells and understanding the role of genes and transcription factors [2, 3]. However, new questions are being asked with the increase in throughput of scRNA-seq and single-cell open-chromatin profiling through ATAC-seq (single-cell assay for Transposase-Accessible Chromatin using sequencing). One such question is, how can we use single-cell transcriptome and epigenome profiles for new applications. Can single-cell epigenome and expression profile help in finding lineage potency of a cell? Can single-cell heterogeneity be used in choosing more specific target pathways for cancer therapeutics? The answers to such questions can be found by representing cell in functional state-spaces. Such as defining cell-state in terms of pathway activity scores could help us to have a meaningful perspective about its role and dynamic behaviour. However, most often enrichment of pathways is done using differential gene expression between two groups of cells and this procedure does not solve the purpose of studying heterogeneity of gene-set enrichment at single-cell resolution.

Another category of methods like SVA[4], RUV[5], scLVM[3] and f-scLVM[6] provide relevance score for known and unknown dominating factors for a group of single-cells. Such methods do not provide enrichment and relevance of gene-sets in each single-cell like PAGODA[7] and AUCell [8]. PAGODA is not designed to handle scRNA-seq data from a non-heterogeneous collection of cells. Whereas, AUCell is used primarily for identification of cells with the activity of chosen gene-sets in scRNA-seq data. Even before PAGODA was proposed, there were a few methods for aggregation of microarray-based expression of genes in gene-sets for downstream analysis procedures [9]. However, when it comes to scRNA-seq profiles, the main hurdle in calculating enrichment of multiple pathways for each single-cell has been the default dependency on read-count data of genes. The read-count values in single-cell profiles are often zero due to true low expression (non-active regions) or dropouts. Dropouts are defined as undetected true expression (activity) due to technical issues. The statistical modelling of read-count of a gene (or genomic site) across multiple cells is a non-trivial task, especially for single-cell open-chromatin and scRNA-seq profiles due to variability in the dropout rate and sequencing depth among cells[7, 10]. Moreover, there has been rarely any attempt to estimate pathway enrichment-scores for single-cells using their open-chromatin profiles for downstream analysis like classification and pseudo-temporal ordering. Hence, there is a need for a uniform method which can transform single-cell expression and open-chromatin profiles from both non-heterogeneous and heterogeneous samples to gene-set activity scores.

In this study, we have addressed the challenge of representing single-cells in terms of pathways and gene-set enrichment-scores estimated using scRNA-seq and open-chromatin profiles despite cell-to-cell variability in dropout of genomic regions and sequencing depth. Unlike previously proposed methods for scRNA-seq profiles, we do not try to normalise or scale read-count of a gene across cells using parametric distributions like Poisson or negative binomial. Scaling read-count across cells with variable dropout rate and sequencing depth increases chances of artefacts. Therefore, we use a common null model to estimate adjusted pathway enrichment scores while handling scRNA-seq profiles. Similarly, while using scATAC-seq profiles, we use the approach of highlighting enhancers by dividing read-counts of genomic sites with their global accessibility scores. We benchmarked our methods and null models for estimating single-cell gene-set enrichment using several published scRNA-seq and scATAC-seq datasets.

Using pathways and gene-set as features for single-cells creates new opportunities and challenges which we tried to explore further. Compared to raw read-counts, the pathway scores of single-cells are more likely to have less sparsity, noise and technical variation, which we exploited for standard procedures like classification and dimension reduction based visualisation. Next, we asked whether the temporal order of cells can be predicted using gene-set enrichment scores as features since it can directly highlight the continuum of lineage potency with inflexion points defined by the activity of pathways. However, existing methods of temporal ordering for single-cell, use read-count or gene-expression matrix which provides less flexibility in terms of avoiding known covariate. Therefore, we developed and included a novel pseudo-temporal ordering method in UniPath which can use pathway scores and allow dropping gene-sets of known covariates. We applied UniPath on a large scRNA-seq data-set of mouse cell atlas (MCA) and performed clustering using pathway scores and annotated many unlabeled cells. We also explored the possibility of using enrichment and co-occurrence (co-enrichment) of pathways for understanding underlying regulation. While analysing the scRNA-seq profile of differentiating hESC cells using pathway scores, we realized the strength of a new way of comparing different populations of cells. Therefore, we performed scRNA-seq for two cell lines of non-small cell lung cancer (NSCLC) and tried to understand the difference in their properties using the new way of comparison.

## Results

For transforming scRNA-seq profiles to pathway scores, we treat each cell separately. Generally, in a single-cell, RPKM (read per Kilobase per million) or FPKM (fragment per Kilo per million) value of genes have a bimodal distribution, where one of the modes is around zero, and other is for non-zero expression values. We used widely and theoretically accepted assumption that most of the time, non-zero RPKM and FPKM values within a sample (or cell) follow log-normal distribution[11]. For a single-cell, we convert non-zero expression values (FPKM, RPKM, TPM, UMI-counts) of genes to P-values (right-tailed) using log-normal distribution (see Methods). UMI-counts do not need to be divided by gene-length to calculate gene-expression values [12]; hence we use log-transformed UMI-counts to calculate P-values. The assumption of log-normal distribution for non-zero gene-expression values for a cell also has support from a report by Furusawa et al. [13] on the ubiquity of log-normal distributions in intra-cellular reaction dynamics. Moreover, skewed distributions often fit log-normal [14], which is also reflected by quantile-quantile(q-q) plots for FPKM, TPM and UMI-counts shown in Figure S1. We apply Brown’s method to combine P-values of genes in a gene-set to reduce the effect of covariation among genes (see Methods). The combined P-value for every gene-set is adjusted using a null background model made using a systematic approach (see Methods, Fig. 1a). The objective of P-value adjustment using a null model created by Monte-Carlo approach (see Methods) is to highlight cell-type-specific gene-set activity and reduce blurring due to background house-keeping function of cells. We call the adjusted P-value of a pathway (or gene-set) in a single-cell as its score.

**Fig. 1:**
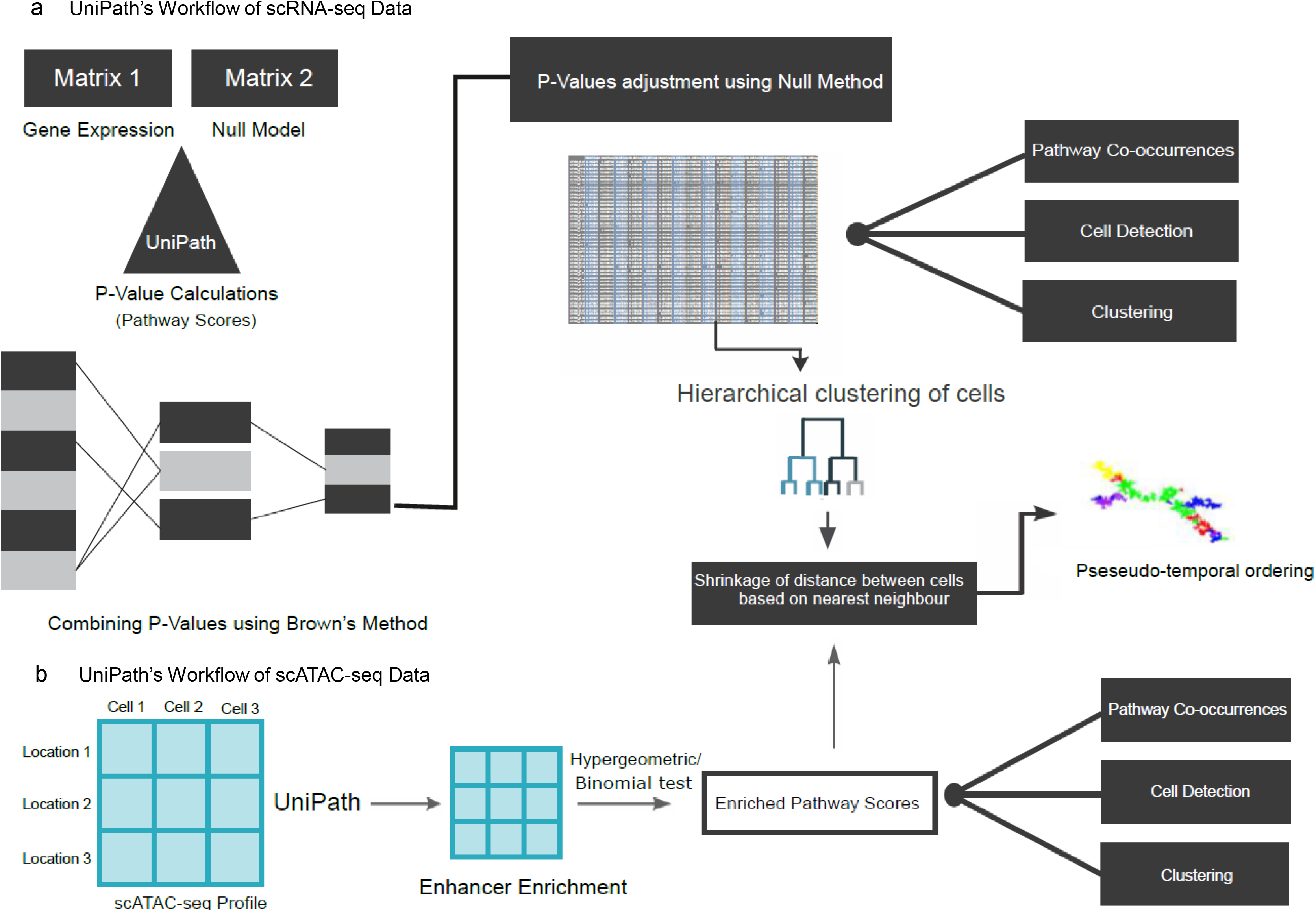
Outline of UniPath. (a) Schematic workflow of UniPath for scRNA-seq data. UniPath works by transforming scRNA-seq gene expression profiles to P-values combined using Brown’s method for each gene-set. The combined P-values are adjusted by using a null model to estimate final pathway score. The null models are made systematically to avoid redundancy. (b) Schematic workflow of UniPath for scATAC-seq data. UniPath transforms open chromatin profiles to pathway enrichment scores for gene-sets by highlighting enhancers and using their proximal genes for Hypergeometric or Binomial test. To highlight enhancers, UniPath normalizes read-count at a peak by its global accessibility score.

### Evaluation of UniPath’s approach of transforming single-cell expression profiles to pathway enrichment scores

Due to the lack of gold standard, it is not trivial to assess gene-set enrichment methods for heterogeneous bulk samples. However, for single-cell from known cell-lines, the marker gene-set for cell-types can be used directly to test methods like UniPath. We used marker gene-sets for cell-types to compare our approach with existing methods PAGODA[7] and AUCell meant for scRNA-seq and GSVA [15]. Systematic evaluation using scRNA-seq profiles from 10 studies (see Table S1) revealed that most of the time UniPath was better than PAGODA, AUCell and GSVA in terms of estimating enrichment of gene-sets for correct cell-type as one among top 5 enriched terms (see supplementary methods). Especially for the non-heterogenous collection of cells, UniPath was substantially better than PAGODA (Fig. 2a, Figure S2, Table S1). Notice that the purpose of using cell-type marker gene-set was to evaluate the correctness of enrichment of gene-sets for downstream analysis steps of clustering and temporal-ordering. To further clarify, we made a collection of gene-sets of non-immune related pathway terms, and as spike-in, we added two known gene-sets related to B cell. We also added two gene-set associated with T cells to the same collection (see Table S2). We checked in what fraction of cells both relevant terms appeared in the top 5 enriched terms (supplementary methods, Fig. 2b). With this control experiment for both B cell[16] and T cell[17], UniPath revealed the correct respective pathways in top 5 enriched terms with substantially better accuracy than PAGODA, AUCell and GSVA (Fig. 2b). To have a further unbiased test, we used GSEA[18] to make reference list of significantly enriched gene-sets (FDR < 0.2) in the group of T cells w.r.t other cell types using mouse cell atlas [19] data-set. We used the reference gene-set as a list of positives for T cell to evaluate UniPath and other three methods in terms of their presence in the top 10 terms in every single T cell (supplementary methods). UniPath had a substantially higher level of presence of reference gene-sets (positives) among 10 term enriched terms in comparison to PAGODA, AUCell and GSVA (Fig2b, Figure S3 a-d). Repeating the same experiments for B cells, we found similar results, highlighting the fact that UniPath is indeed superior in estimating cell-type-specific enrichment for gene-sets in single cells.

**Fig. 2:**
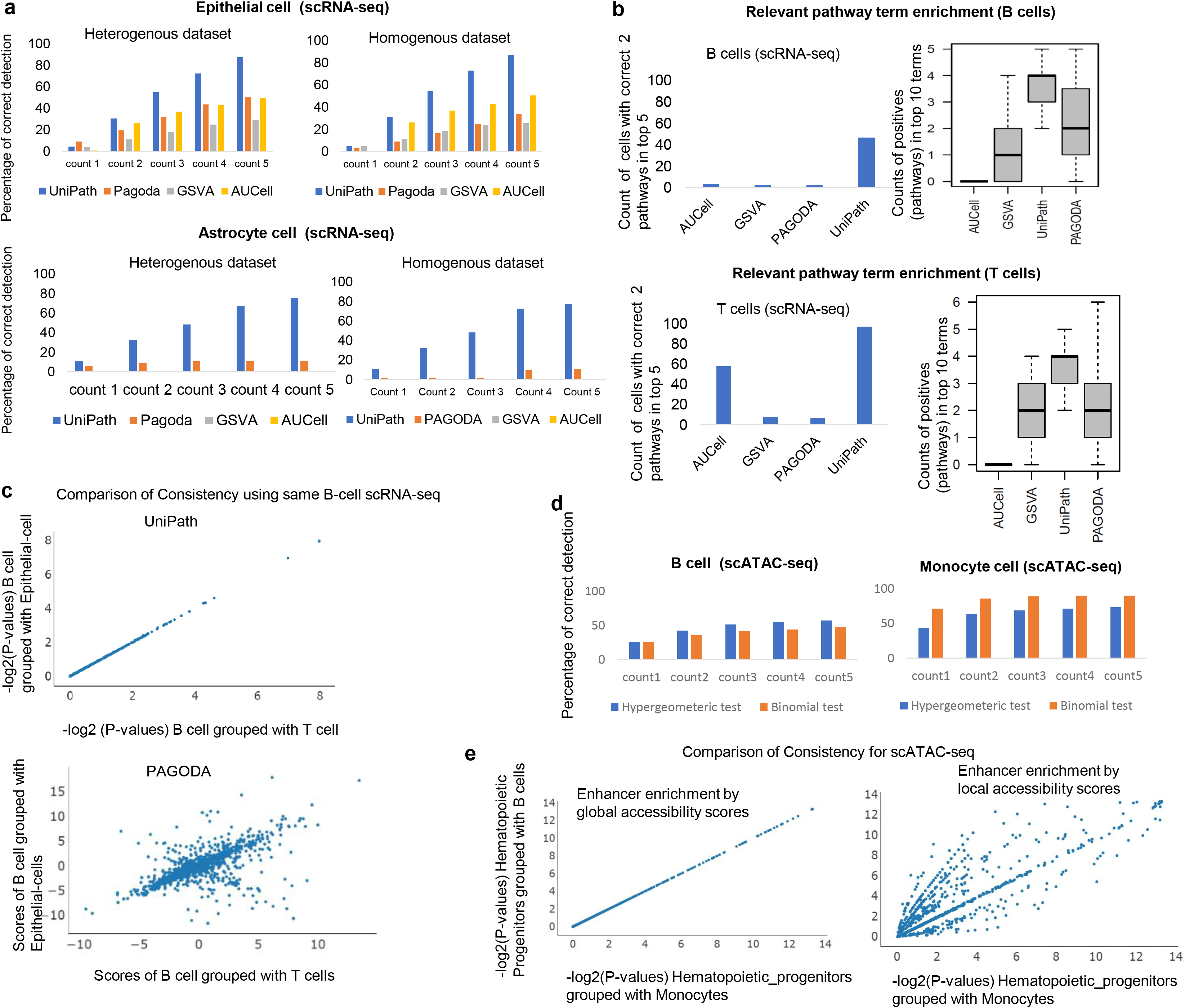
Evaluation of UniPath using scRNA-seq and scATACseq profiles. (a) Accuracy of highlighting relevant gene-set among top enriched terms. The terms here are cell-types, and gene-sets are set of marker genes for cell-types. The results for scRNA-seq profiles of Epithelial cells and Astrocytes, as shown here. The evaluation was performed using both homogeneous and non-homogeneous data-sets (Table S1). More such examples are shown in Figure S2. (b) Accuracy of results of pathway enrichment by UniPath and other 3 methods for B and T cell scRNA-seq profiles. For systematic evaluation, two figures are shown corresponding to two lists of gene-set. Left figure is made using a list consisting of non-immune pathways and two pathways (positives) relevant to T cell and other two gene-set for B cells (bar-plot). The figure on the right (box-plot) was made using a list of all gene-sets, but positives decided based on GSEA applied to relevant cell types (T or B cell) in mouse cell atlas. The count of positives in top 10 enriched terms in every cell is shown. (c) Consistency of UniPath based pathway-scores when B cells are grouped with epithelial or T cells in comparison to PAGODA. Gene-set enrichment scores provided by PAGODA change when the same cell is grouped with other cells. While UniPath’s output remains consistent. (d) Evaluation of UniPath for highlighting correct gene-sets among top enriched term for single-cell ATAC-seq profile. For this purpose, marker gene-sets for cell-type were used by UniPath on the scATAC-seq profile of B cell (GM12878) and Monocyte. Here UniPath used global accessibility scores to highlight enhancers. (e) Consistency of UniPath based pathway enrichment score calculation using scATAC-seq. Here hematopoietic progenitor cells are grouped with B cells or monocytes. UniPath’s approach of highlighting enhancers by using global accessibility score gives a more consistent result than mean based normalisation (local accessibility score).

We also assessed the consistency of enrichment of pathways by UniPath and the other three methods (PAGODA, AUCell, GSVA). We analyzed B cells (GM12878) scRNA-seq profile [20] while grouping them each time with different cell types. The scores for pathways and gene-sets from PAGODA and GSVA were not consistent (see Fig. 2c, Figure S3e) and for every cell, the output was dependent on the composition of cell type in the data-set. However, UniPath and AUCell based enrichment scores for a cell remains consistent and is not affected by other neighbouring cells (Fig. 2c, Figure S3e). Thus, UniPath also resolves the issues of highlighting correct gene-sets and relevant pathways with consistency for each single-cell irrespective of the level of heterogeneity of cell-types in the provided scRNA-seq data.

### Gene-set enrichment with UniPath as an alternative dimension-reduction method for single-cell ATAC-seq profile

For the transformation of open chromatin profile of single-cells to pathway enrichment scores, UniPath first highlights enhancers by normalizing read-count on peaks using their global accessibility scores (Fig. 1b) (see Methods). For this purpose, it intersects peak list of given scATAC-seq profile with a pre-compiled list of genomic regions with pre-calculated global accessibility scores. The global accessibility score of a genomic site is proportional to the number of times it appeared as a peak in available bulk open-chromatin profiles. (see Methods). The motivation behind normalising read-count of each peak using its global accessibility score is to have consistency and avoid adjustment of variability in sequencing depth and dropout rate. For every cell, genomic sites with high normalized read-count are chosen as foreground set. Then, for every cell, UniPath uses proximal genes of peaks in its foreground set to estimate statistical significance (P-value) of enrichment gene-sets using Hypergeometric or Binomial test. We call the P-value of enrichment of a pathway or gene-set as its score. We performed systematic evaluation using cell-type marker gene-set for both bulk ATAC-seq of immune cells[21] and multiple single-cell ATAC-seq profiles [22, 23]. Most of the time, UniPath highlighted correct cell type among top 5 enriched gene-set for both bulk and single-cell ATAC-seq profiles (Fig. 2d, Figures S4 and S5). Making a global list of peaks with accessibility score is possible due to the availability of bulk open-chromatin profiles. In the absence of enough publicly available open-chromatin profiles for any species, one can also use UniPath by calculating local accessibility score (study-specific normalisation). However, local accessibility scores are dependent on the composition of cells in the data-set and could lead to inconsistency in estimation of enrichment of gene-sets (shown in Fig. 2e). Thus, UniPath calculates consistent and mostly correct enrichment scores for pathway and gene-sets for every cell using its scATAC-seq profile independently.

### Handling dropout and batch effect and evaluation using visualisation and clustering

Most often in single-cell scRNA-seq profile, there is heterogeneity in dropout rate among cells. Such dropout of genes could be random or systematic. The systematic dropout usually occurs due to differences in sequencing depth or RNA degradation level (frozen vs fresh) among different batches of samples. We tested whether UniPath is robust to systematic dropout variability among cells. We simulated the systematic dropout rate using publicly available scRNA-seq data-sets [24, 25] (Fig. 3a) (see Methods). We found that applying PCA on raw read-count lead to artefactual cluster formation due to systematic dropout. Whereas, using UniPath based pathway scores, similar cells remained in the same cluster irrespective of the non-random pattern in dropout rate (Fig. 3a, Figure S6a). Besides being robust to systematic dropout, UniPath allows correction for batch effect before calculating the adjusted p-value for the enrichment of pathways (see methods and Figure S6b). The framework of UniPath avoids normalisation artefact due to sequencing depth and dropout rate; therefore, it could be used for efficient classification of single-cell. During hierarchical-clustering, UniPath based gene-set scores provided comparable or higher clustering-purity than raw expression based results [20] (Figure S6c-d). We further compared visualisation and clustering using pathway scores from 4 different methods for scRNA-seq profiles. We used t-SNE based visualisation and performed dbscan[26] based clustering of t-SNE coordinates (supplementary methods). We tried using different values for eps parameter (for the radius of the neighbourhood) in dbscan. However, for almost all values of eps parameter of dbscan, UniPath based pathway scores provided better clustering purity. UniPath based clustering results also had higher values for adjusted rand Index (ARI) and normalised mutual information (NMI) than other three methods (PAGODA, AUCell, GSVA) (Fig. 3b and Figure S7). Due to better visualisation and clustering we found meaningful sub-groups of stromal cells in an organ (Uterus) in mouse cell atlas only while using UniPath scores (see Figure S7c and supplementary information).

**Fig. 3:**
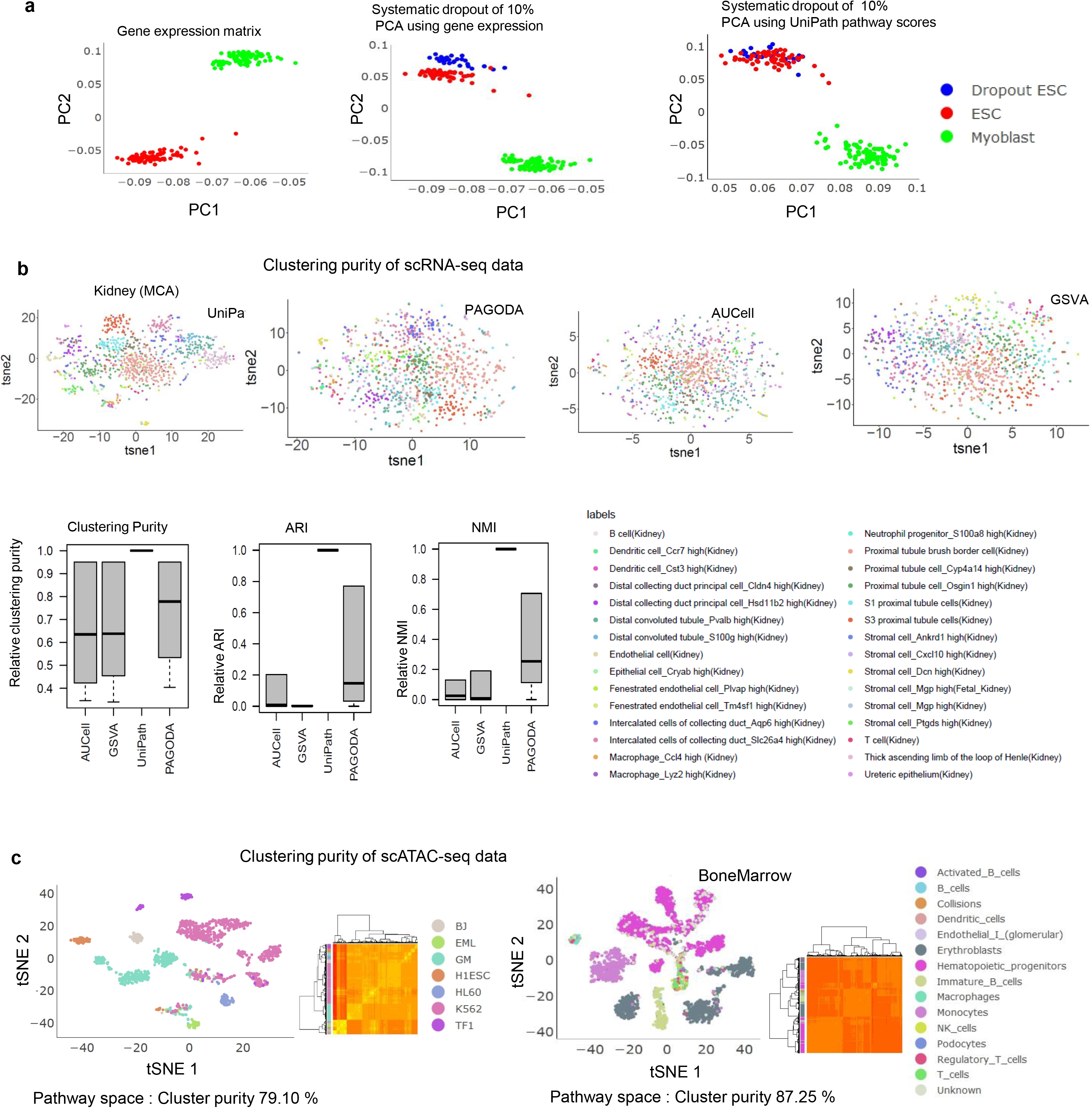
Reduction of the artefact by Unipath and clustering using pathway scores. (a) Principal component analysis (PCA) based visualization of data consisting of human embryonic stem cells (hESC) and Myoblast cells scRNA-seq profile. Here PCA was done using scRNA-seq based gene expression. Simulation of systematic dropout of 10% genes in few hESCs leads to the formation of a separate group of hESCs in PCA based visualisation of raw gene expression. However, PCA based visualisation using pathway score from UniPath shows a grouping of all ESCs in same cluster irrespective of systematic dropout. (b) t-SNE based visualisation of scRNA-seq profiles after transformation to pathway scores for cells in the kidney from mouse cell atlas. Here UMI-counts were used for calculating gene-set scores. The boxplot show efficiency of clustering at different values of “eps” parameter of dbscan method. Three types of clustering efficiency quantification (clustering purity, adjusted rand index and normalised mutual information) are shown here. The clustering efficiency is shown relative to the corresponding value for UniPath based results as relative-ARI, relative-NMI at different values of ‘eps’ parameter in dbscan method. (c) Clustering purity of scATAC-seq profiles transformed into pathway space. Scatter plots of tSNE coordinates for cells from two scATAC-seq datasets (Cusanovish et al. [59] and Buenrostro et al.[22]) are shown here.

When hierarchical clustering was performed using pathway scores of imputed scATAC-seq profiles, it resulted in high clustering-purity (Fig. 3c). We also compared visualisation and clustering of scATAC-seq profiles with the outputs of two other tools (ChromVar and SCALE) [27] [28] which are used for handling scATAC-seq count but not for finding enrichment of gene-set. UniPath based pathways scores for imputed scATAC-seq profiles provided better or comparable visualisation and clustering when compared to the output of ChromVar and SCALE (Figure S8). Thus high accuracy in clustering with gene-set enrichment scores, proves that defining cell-states in terms of pathway enrichment using UniPath can be a reliable method for classifying single-cell epigenome and transcriptome data-sets.

### Pseudo-temporal ordering using pathway enrichment scores and visualizing the continuum of lineage potency and pathway co-occurrence

Pathway-scores based representation can provide new similarity measures among cells as well as help in avoiding few covariates like cell-cycle phase, tissue-microenvironment or culture conditions. However, current methods for temporal ordering [29] of cells are designed to handle FPKM and read-counts of genes, and they are also not meant for visualisation of the continuum of pathway scores on temporally-ordered cells. Hence, we extended UniPath with a novel method for the pseudo-temporal ordering of single-cells which can utilize pathway scores based representations. For temporal ordering, we apply two levels of shrinking of distances between cells based on their pre-classification and continuum among their classes before finding a minimum spanning tree (MST). To find a continuum between different classes, we use KNN based approach after initial classification so that correct temporal ordering among clusters of cells can be determined (see methods). Without shrinkage of distances using pre-classification and KNN based continuum, the MST could not capture correct temporal order in few cases (see Figure S9 a-b). Overall, we found that UniPath is indeed able to predict approximately correct order of cells using pathway scores derived from scATAC-seq and scRNA-seq profiles (see Fig. 4 and Figure S9).

**Fig. 4:**
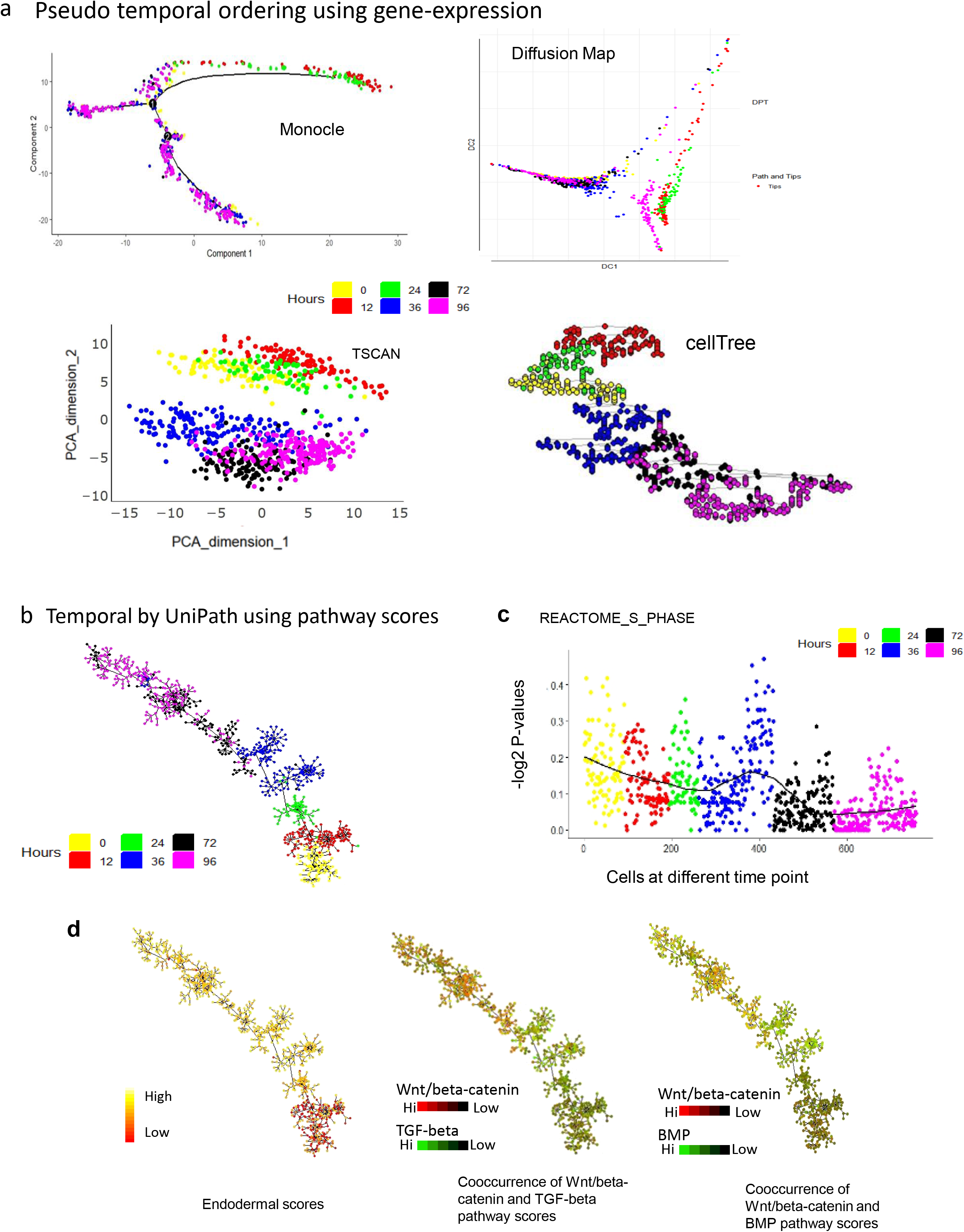
Pseudotemporal ordering using gene set enrichment scores and visualisation of potency and pathway co-occurrences. The dataset used here, consisted of cells collected at different time points (0,12, 24, 36, 72 and 96 hours) of differentiation of human embryonic cells (hESC, 0 hours) towards definitive endoderm (DE) [24] (96 hours). (a) Imperfect prediction of temporal order using gene-expression by other tools. Monocle mixed 0 hours (hESCs) and 96 DE hours cells, Diffusion map also mixed 0 hour cells with 72 hours. TSCAN could not find a proper order in the sequence of 0, 12, 24, 36, 72 and 96 hours, cellTree also could not find a proper temporal order among cells (b) Predicted temporal order of cells of differentiating human embryonic stem cells towards definitive endoderm. The order predicted is exactly according to true time-points of cells. (c) The enrichment score of gene-set for S phase in cells at different time points of differentiation. (d) The trend of endoderm lineage potency and co-occurrence of pathways at single-cell resolution on temporally ordered tree.

We further used UniPath for temporal ordering of scRNA-seq profile of Human embryonic stem cells (hESC) and their differentiated states collected at time points of 0, 12, 36, 72 and 96 hours during differentiation towards definitive endoderm (DE) (Fig. 4)[24]. Using other tools (monocle, TSCAN, DiffusionMap, CellTree)[25, 30–32] for pseudo-temporal ordering with gene-expression (Transcript per million, TPM) matrix (Fig. 4a), resulted in predicting wrong order of cells for the same data-set. However, with UniPath, when we dropped gene-set associated with cell cycle, we achieved the correct order of cells. We found that the score of gene-set for cell cycle (S phase is shown here) is higher at 0 and 12 hours, possibly due to a high level of proliferation (Fig. 4c). The S phase gene-set score kept decreasing as the cells differentiated towards endoderm (Fig. 4c). However, at 36 hours, we find two batches of cells such that one batch of cells had a much lower score of cell-cycle (S phase here) gene-set than the other. Such batches of cells hint about the possible impact of cell-cycle as a covariate during the prediction of temporal order. Besides handling known covariates, UniPath can also be used to visualise continuum of lineage potency and concurrence of two pathways on pseudo-temporally ordered tree. The endodermal lineage gene-set score increases as the cells differentiate towards endoderm (Fig. 4d) (see methods). Such as Wnt/beta-catenin and BMP pathway scores seem to have mild co-enrichment at 0 and 12 hours. As cells differentiated towards the mesendodermal stage at 24 and 36 hours, the BMP signalling pathway seems to be getting more enriched compared to WNT/beta-catenin. Importance of BMP till mesendoderm stage had been shown before [33]. After 36 hours enrichment level of WNT/beta-catenin is slightly higher; however, its co-occurrence with BMP shows a slow increase towards the end of temporally ordered tree (Fig. 4d). UniPath also enables analysis of co-occurrence pattern and detection of clusters of pathways which can be used to infer context-specific regulation (Figure S10, Figure S11, supplementary information). Overall, UniPath tends to be beneficial for predicting correct temporal order of cells and making inference about stage-specific co-occurrence of pathways during differentiation of cells.

### Enabling analysis of large atlas scale scRNA-seq data-set using pathway enrichment scores

The consistency due to the use of global null models by UniPath provides horizontal scalability in calculating scores for pathways for single-cells. Even with a single CPU, the computation time needed by UniPath is much less than PAGODA on the same number of cells (Fig. 5a). The horizontal scalability, speed and consistency of UniPath allowed us to transform expression profiles (UMI-counts) of more than 61000 single-cells from mouse cell atlas (MCA) dataset[19], by dividing them into smaller groups of cells. Such step of the division of data-set and transformation to pathway scores with other similar tools (PAGODA) would provide inconsistent results, as explained and demonstrated above. We selected 49507 cells which have more than 800 genes with non-zero expression value (see supplementary methods). Further, t-SNE [34] based dimension reduction of pathway scores and subsequent application of dbscan[26] (Additional file 1), revealed a correct grouping of most of the cells according to their tissue (Fig. 5b, Figure S12). As expected, some cells did not group with their source tissue cluster, but they formed a separate class. Such as immune cells from different organs grouped together in cluster 13,14,15 (see Fig. 5b, Additional file1).

**Fig. 5:**
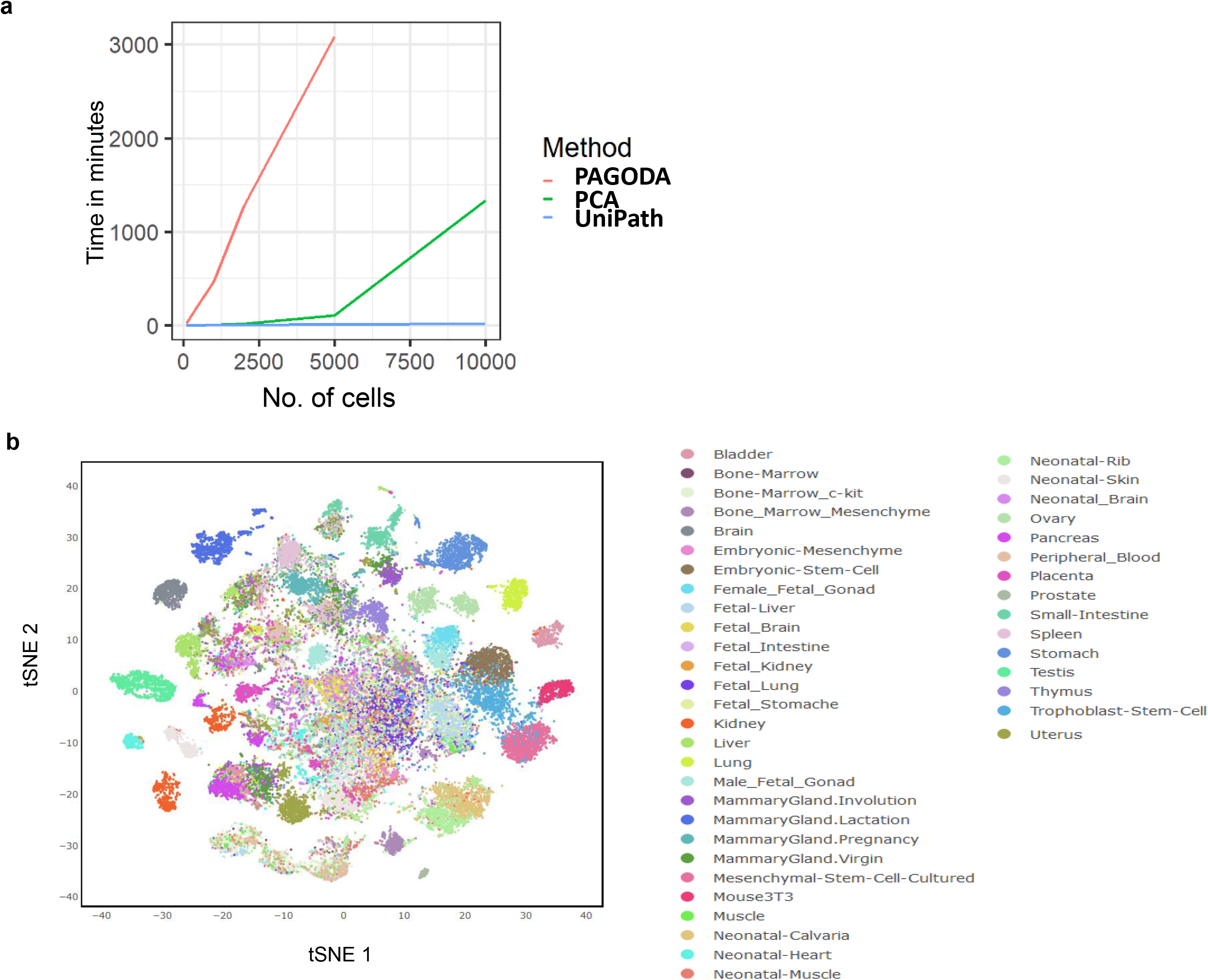
Execution time of UnPath and analysis atlas scale single-cell RNA-seq data-set. (a) Comparison of the execution time of UniPath with PAGODA and PCA with varying number of cells. (b) Scatter plot of t-SNE results for 49507 cells of mouse cell atlas (MCA) data-set [19] represented using pathway scores. The transformation of MCA data-set to pathway enrichment scores was possible due to consistency and scalability provided by UniPath. The clusters detected using the shown tSNE result for MCA data-set are shown in Figure S12. For MCA data-set UMI-counts were used by UniPath to calculate pathway scores.

Surprisingly, co-clustering of few non-immune cells from different tissues revealed convergence which has been rarely reported before by single-cell analysis but is supported by earlier scientific studies. Such as in our analysis, cluster 40 has *Afp+* fetal liver hepatocytes as well as *Afp+* placental endodermal cells which were reported to belong to different classes in the original study of MCA (Additional file 2 and 3). Cluster 40 also has a few Fabp1+ hepatocytes. It has been previously shown, that placenta-derived multipotent cells (PDMCs) with the expression of Afp (see Fig. 6a) gene, has endodermal features and can differentiate easily towards hepatocytes like cells[35]. We compared both types of cells (*Afp+* placental endodermal cells and *Afp+* fetal liver hepatocyte) in cluster 40 with other cells in MCA. Among top 50 pathways more enriched in *Afp+* placental endodermal cells (from cluster 40) 22 were also present in 50 most differentially upregulated pathways in hepatocytes cells of cluster 40. These common 22 pathways (44% overlap) were mostly related to lipid metabolism (see Table S3). However, there was certainly a difference between hepatocytes and *Afp+* placental endodermal cells of cluster 40 in t-SNE based visualization. (Fig. 6a).

**Fig. 6:**
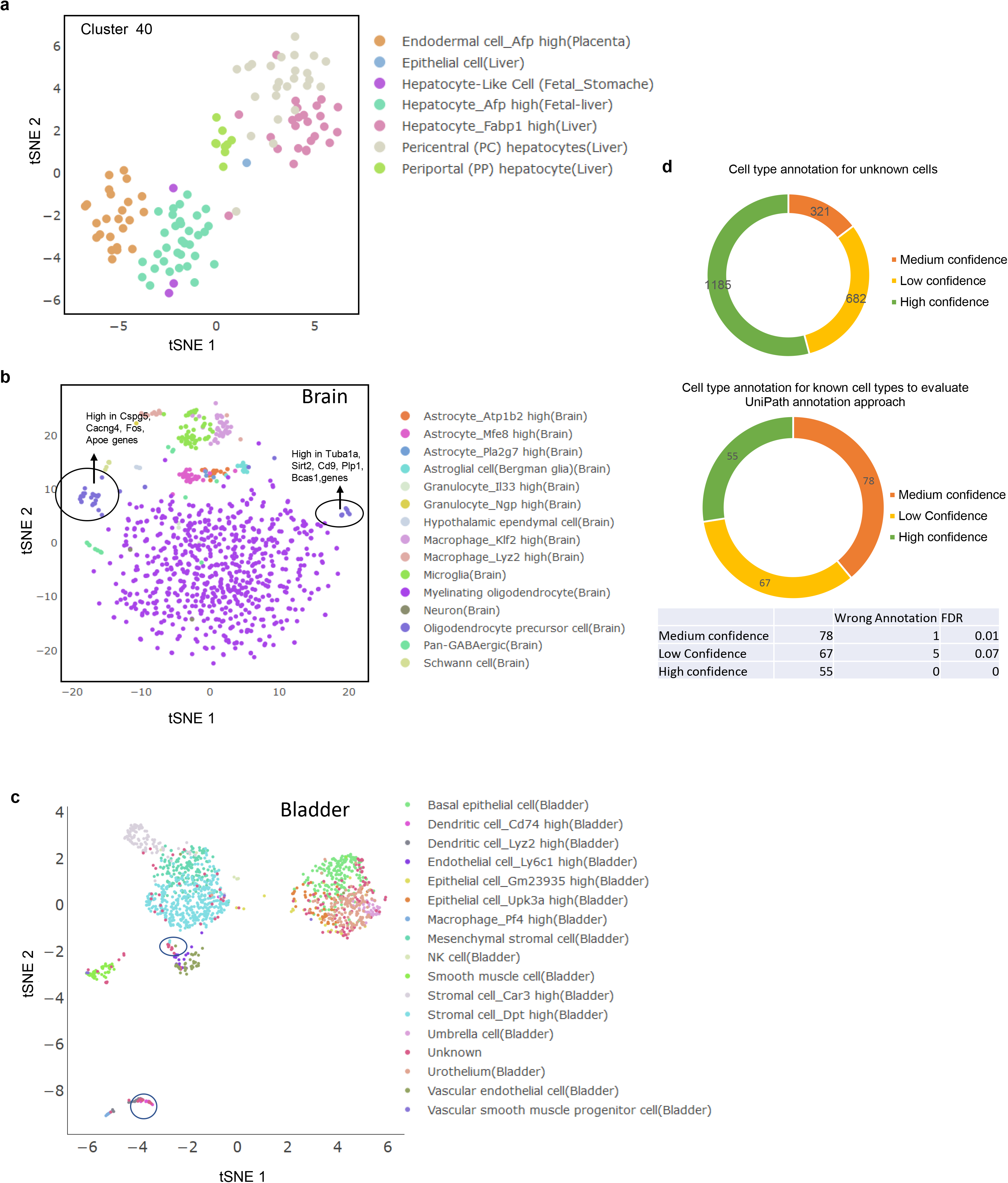
Analysis of single-cell RNA-seq profile of Mouse cell Atlas (MCA) (a) t-SNE scatter plot of cells co-clustering in cluster number 40. AFP high placenta endodermal cells and Afp+ hepatocytes do not overlap, but they lie closer to each other in t-SNE based plot. (b) t-SNE results of the scRNA-seq profile from brain showing two clusters of oligodendrocyte precursor cells along with their enriched genes. These two clusters of cells were labelled as a single cell-type in the original study by Han et al. [19] (c) t-SNE based scatter plot for bladder cells in MCA data-set represented in terms of pathway enrichment scores. Two clusters of cells labelled as “unknown” were visible. Cells in one of the two clusters were identified as cd74_high_dendritic cells. (d) Pie chart showing confidence level for cell-type annotation on MCA data. Total 2188 cells with “unknown” label could be annotated with the help of UniPath. False detection rates for different confidence level are also shown for the same procedure when the same procedure was applied to 200 randomly chosen but labelled cells.

Another example of convergence is cluster 3, which has virgin mammary gland luminal-epithelial cells (including alveoli cells) and glandular epithelial cells from uterus. An interesting example of convergence is cluster 52, that has *Col10a1+* and *Cmnd+* bonemarrow mesenchyme stromal, pre-osteoblast and chondrocytes cells (Additional file 2). It is well-known that bonemarrow mesenchyme stromal (also known as mesenchymal stem cells [36]) has a high potency to transform to pre-osteoblast and chondrocytes cell state[37]. In contrast to such a result, *Cxcl1+* MSC from in vitro culture grouped with trophoblast stem cells in cluster 21. It can be noticed, that the cell types from different organs, converging in a major cluster, did not overlap completely with each other but formed their own sub-cluster within their major class (Fig. 6a). However, the convergence to a major class shows a reduction of covariates due to underlying tissue microenvironment in gene-set scores, which caused cells with a similar state to group together. Overall UniPath, provided a new dimension to classify cells and revealed that even though an organ has a specific type of cells for its functioning, it also has some cells with a regulatory state similar to cell-types from other parts of the body.

#### Revealing new minor classes using pathway scores and annotation of unlabeled cells

When an analysis is performed using an expression-matrix of a large number of genes, feature selection is often needed for proper clustering. However, currently, there is no optimal solution for selecting genes to highlight all relevant classes. Feature extraction in terms of pathway scores can help to reduce noise, sparsity and effect of few covariates. Thus, pathway scores can help to highlight clusters of cells which might not get detected by using raw read-counts. Such as analysis using pathway scores of brain cells in MCA data-set, resulted in the detection of a new cluster among oligodendrocyte-precursor cells. Oligodendrocyte precursor cells belonging to the new small cluster had a higher expression for *Tuba1a, Sirt2, Cd9, Plp1* and *Bcas1* (Fig. 6b). These genes are involved in the differentiation of oligodendrocyte-precursor towards mature oligodendrocytes[38–41]. On the same trend, we found two new clusters of “unknown” cells from the bladder in MCA data-set (Fig. 6c). We could annotate cells in one of the newly detected clusters in the bladder as Cd74_high dendritic cells.

Despite a tremendous effort by Han et al. [19] they could not annotate all cells in their MCA dataset, hence among 49507 cells, 5590 cells had annotation as “unknown”. We could find cell-type for 2188 cell with “unknown” label, using a two-pronged approach enabled by UniPath (see Fig. 6d, supplementary methods, Additional file 4). Our approach is two-pronged as it utilizes UniPath scores for cell-type marker-set as well as the result of sub-clustering (supplementary method). When we used the same procedure on 200 randomly picked cells with labels, we achieved a false detection rate (see Fig. 6d) of less than 8% even for low-confidence annotation.

### Application in inferring context-specific regulation in cancer cells

We further explored how UniPath can help in for studying context-specific regulation in cancer cells which is often required for precision oncology. Recently Wang et al. [42] showed a difference in metabolic profile among two types of NSCLC cell lines, non-adherent tumorspheres (TS) grown in serum-free culture conditions and adherent (Adh) cells cultured in serum-containing medium. They have demonstrated a high level of the tumorigenic potential of non-adherent TS cells in comparison to adherent ones using mouse xenograft models. We performed single-cell expression profiling of 162 cells out of which approximately half were TS cells, and others were Adh cells (see Methods). We asked whether we can utilise the difference between TS and Adh cells in highlighting signalling pathways associated with tumorigenicity in lung cancer. After applying UniPath, the differential enrichment analysis using Wilcoxon Rank sum test revealed GPCR ligand binding gene-set (Fig. 7a, Figure S13a), IL23 pathway, cytochrome_P450_drug_metabolism and prolactin_receptor_signalling as having higher enrichment in TS cells (based on median fold change and Wilcoxon Rank-sum P-value < 0.01, Fig. 7a) (Additional File 5). Distributions and gradient of some pathways in TS and Adh cells are shown in Figure S13. GPCR and IL23 signalling are known to be associated with plasticity and proliferation of NSCLC[43–45]. Cytochrome P450 is also involved in promoting tumour development [46].

**Fig. 7:**
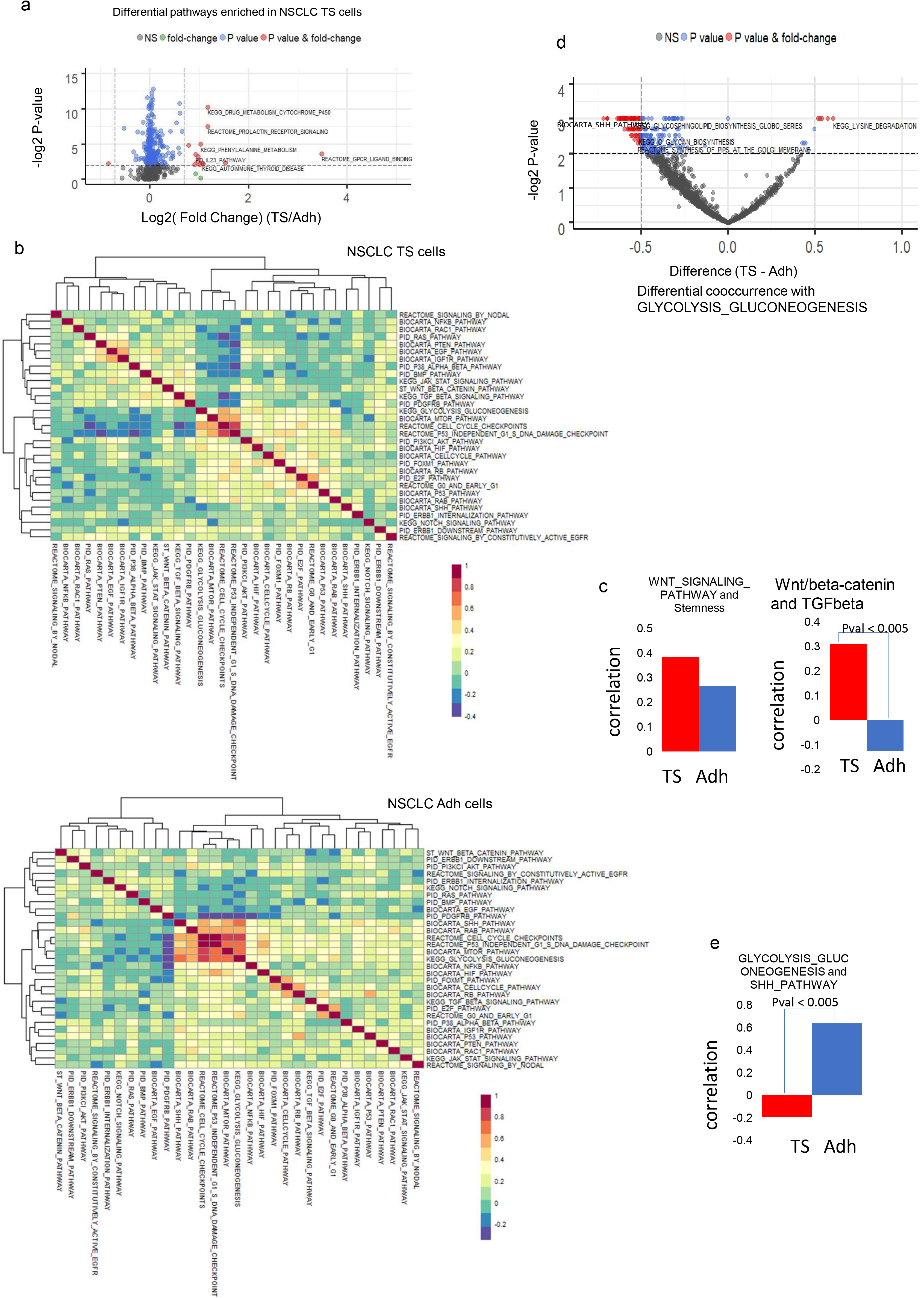
Differences in enrichment and co-occurrence of pathways in two types of cells of non-small cell lung cancer (NSCLC). FPKM values were used for estimating gene-set enrichment scores by UniPath. a) A global view of differential enrichment of pathways using volcano plot. The x-axis shows log fold-change of median enrichment scores of gene-sets in tumorosphere (TS) and Adherent cells (Adh). P-values were calculated using the Wilcoxon rank-sum test. Few pathways which showed a significant difference in enrichment between TS and Adh cells are shown here. (b) Heatmaps of correlation between pathways are shown with their hierarchical cluster in TS and Adh lung cancer cells. (c)Correlation values of WNT signalling pathway with gene-set of stemness in TS and Adh cells are shown as a bar plot. The other bar-plot shows co-occurrence of WNT/beta-catenin with TGF-beta signalling pathways in Adh and TS cell lines. WNT/beta-catenin and TGF-beta had significant differential co-occurrence among TS and Adh. (P-value < 0.005) (d) Volcano plot showing differential co-occurrence of pathways with gene-set for Glycolysis_Gluconeogenesis among TS and Adh cells. Sonic Hedgehog (SHH) pathway and glycolysis gene-set had significant differential co-occurrence among TS and Adh cells. (e) Spearman correlation of scores of Gycolysis_Gluconegenesis pathway with sonic hedgehog pathway (SHH) pathway in TS and Adh cells. The P-value of differential co-occurrence is also shown.

We further used an approach, rarely used for scRNA-seq. We performed co-occurrence and differential co-occurrence analysis for pathway and gene-set pairs. Wnt pathway had the highest correlation with stemness gene-set in TS cells. However, in Adh cells, Wnt was not among top correlated pathways with stemness gene-set. We found that Wnt/beta-catenin pathways had a significantly higher correlation with the TGF-beta pathway in TS in comparison to Adh cells (P-value < 0.005, Jaccard index=0, see Table S4). However, TGF-beta pathway itself did not have a significant difference in enrichment among TS and Adh cells (Figure S13a). Both Wnt/beta-catenin and TGF-beta are known to promote the state of epithelial to mesenchymal transition (EMT) in cancer cells which is associated with high tumorigenicity [47]. Moreover, it has been previously shown that simultaneous over-activation of Wnt/beta-catenin and TGF-beta signalling promotes tumorigenicity and chemo-resistance in NSCLC cells [48]. Using hierarchical clustering of 31 chosen pathways, we found that TGF-beta, Wnt/beta-catenin and PDGFRB pathways co-clustered together in TS cells. In contrast, in Adh cells WNT/beta-catenin pathway grouped with ERBB1 and PI3K1 signalling. The difference in co-occurrence pattern of Wnt/beta-catenin pathway in TS and Adh cells (Fig. 7 b-c) and prior knowledge about the effect of their co-stimulation with TGF-beta in NSCLC hints about a possible cause of higher tumorogenicity in TS cells.

Wang et al. [42] also reported that glycolytic intermediates are more enriched in Adh cells. Our analysis revealed that among non-metabolic gene-sets, sonic hedgehog (SHH) pathway had the highest level of differential co-occurrence (P-value < 0.005, Jaccard index=0) with glycolysis gene-set (Fig. 7d). SHH and glycolysis pathway had a correlation of 0.63 in Adh cell compared to −0.02 in TS cells (Fig. 7e). SHH pathway is known to be promoting glycolysis in multiple types of cancer[49]. In our hierarchical clustering result (Fig. 7b), SHH pathway also seems to group with cell-cycle related gene-sets which hints about its involvement in the regulation of proliferation in Adh cells. Previously SHH pathway has been associated with proliferation and drug-resistance in NSCLC[50]. However, our analysis reveals that its role is context-specific, and it could have a more dominating role in Adh like NSCLC cells compared to TS cells. Similarly, many more such differences could be revealed among Adh and TS cells. However, our analysis here is meant to show that UniPath can help in building relevant hypothesis and help researchers in designing follow-up study of context-specific regulation in cancer cells.

## Discussion

Exploiting single-cell heterogeneity using pathways and gene-set enrichment can give rise to multiple new applications. However, it needs an estimation of consistent enrichment scores for gene-set. UniPath fills the gap between the demand for consistent gene-set enrichment scores for a multitude of applications and the availability of single-cell transcriptome and open-chromatin profiles. The novel approach of processing each cell independently, using a global null model provides consistency and scalability to UniPath for calculating gene-set enrichment. Such approach of UniPath also provided independence of normalization w.r.t other cells which is needed to handle cell-to-cell variability in noise, dropout rate and sequencing depth. UniPath is robust to systematic dropout as well as it can handle batch effect in scRNA-seq profiles. Compared to other similar methods (PAGODA, AUCell and GSVA), UniPath based pathway scores provided better visualisation and classification accuracy for both UMI and non-UMI based scRNA-seq profiles. Thus, UniPath’s approach of approximating the distribution of UMI-count of scRNA-seq to log-normal can provide a satisfactory result. In the absence of a consensus about the distribution of non-zero UMI-count within a cell, we would use lognormal distribution for UMI-count, but we may update our approach in future.

For both scRNA-seq and scATAC-seq profiles, there is a similarity in the downstream procedures after the transformation to gene-set enrichment score. Thus, UniPath provides a uniform platform for analyzing both single-cell transcriptome and open-chromatin profiles with the new dimension of pathway enrichment scores. Especially, for scATAC-seq profiles, we have shown for the first time that transformation to pathway enrichment scores does not reduce the purity of classification of cells. UniPath also provides an alternative solution to transform more than one scATAC-seq read-count matrices to same feature space, despite differences in their peak list. In addition, we have shown how UniPath can help to use pathway scores for temporal ordering and displaying co-enrichment pattern among them.

Due to its horizontal scalability and consistency, UniPath helped in analysis of large MCA scRNA-seq data-set (61000 cells). Classification of MCA data-set using pathway scores revealed few clusters in which one of its member cell-type could be easily differentiated to the other. Such as cluster 40 having *Afp+* placental endodermal cells and fetal liver hepatocyte[35] and cluster 52 with bonemarrow mesenchyme stromal, pre-osteoblast and chondrocytes cells[37]. It could be the result of new regulatory distance defined by pathway scores and suppression of covariate due to tissue micro-environment. Such results hint that biologist could use UniPath to find convergence and feasibility of convertibility between different cell-types.

There is a vast literature on the non-trivial problem of analysing patterns of enrichment and co-occurrence of pathways using bulk expression profiles[51–53]. Exploiting heterogeneity among single-cells with UniPath can easily leverage such analysis. UniPath can be used to elucidate three kinds of differences between two populations of single cells. First one is differential enrichment of pathways, that is used quite often. The other two possibilities with UniPath, which have been rarely explored with single-cell scRNA-seq and scATAC-seq, are analysis of modules (clusters) of pathways and enumeration of differential co-occurrence between two pathways. Such as, it revealed an increase in enrichment of Nodal signalling with differentiation of hESC towards DE and patterns of its co-occurrence with other pathways (SMAD2, Wnt/beta-catenin) which are corroborative with existing literature (see supplementary Methods, Figure S10). Similarly, UniPath allowed us to study pathway co-occurrence in two types of NSCLC and differential co-occurrences of few pathway pairs (TGF-beta and Wnt/beta-catenin; SHH and glycolysis) which had literature support. Hence, besides improving classical procedures like classification and cell-type detection, UniPath also open avenues for new kinds of analysis for scRNA-seq and scATAC-seq profile.

## Conclusion

We have developed methods for representing single-cells in terms of pathway and gene-set enrichment score. Due to the approach of using a global null model for single-cell expression profile, the result of our method is more accurate and consistent than the existing tools. For single-cell ATAC-seq profile, the approach of using global accessibility score to highlight enhancers has made our method robust and consistent in estimating pathway enrichment scores. We have also supplemented our methods with a unique function of determining the temporal order of cells using their pathway enrichment scores. Using pathway scores as features for temporal ordering provide the benefit of avoiding covariates by dropping chosen gene-sets. Having a consistent and robust estimate of pathways and gene-sets enrichment scores opens avenues for multiple other kinds of analysis which we explored. Such as gene-set enrichment scores 1) define new kind of regulatory distance among every pair of cells 2) enable estimating differential enrichment of pathway among two populations of cells 3) allow estimating co-occurrence and differential co-occurrence among two pathways and 4) finding a cluster of co-regulated pathways. For mouse cell atlas data-set, using new kind of regulatory distance defined by pathway scores revealed biologically meaningful co-clustering among cells from distal organs and highlighted previously undetected cell-clusters and helped in annotating unlabeled cells. Using our novel single-cell RNA-seq data-set of two types of lung cancer cells and analysis of pathway clusters and co-occurrences revealed the potential of UniPath for building relevant hypothesis about context-specific regulations in cancer cells.

## Methods

### Calculating Enrichment of gene-sets for scATAC-seq profiles

Multiple kinds of regulatory sites like promoters, enhancers and insulators have higher chromatin accessibility than background genomic regions. Most of these regulatory sites like insulators and active promoters tend to have high chromatin accessibility in the majority of cell types. However, to estimate differences among single-cells using open chromatin profiles, sites with cell-type-specific activity like enhancers could be more useful. Moreover, the profile of enhancers provides a more clear perspective about active pathways in a cell. Therefore, UniPath first normalizes the tag count of scATAC-seq profiles of each cell to highlight enhancers. It has two methods for normalization to highlight enhancers. In the default first method, normalization is done using precalculated global accessibility score of genomic sites. For multiple organisms like human, mouse and Drosophila, bulk sample chromatin accessibility is available for many tissues and cell types. A union list of open-chromatin sites was made for human hg19 version genome, and the accessibility score of a union list of sites was calculated. For example, for Human hg19 genome, we combined DNAse-seq and ATACseq peaks from ENCODE and IHEC consortiums[54], to achieve more than 1 million sites and calculated accessibility scores of combined peak list (see supplementary methods). The accessibility score is calculated for a site as the proportion of cell types or samples in which it was detected as an open-chromatin peak.

For tag-count p_ij_ of a peak i in a single-cell j, the normalisation is done as

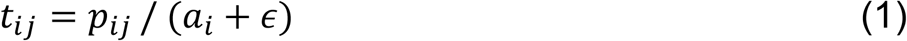

where ϵ stands for a pseudo-count and *a*_*i*_ is the global accessibility score for peak *i*. Thus, the first method of highlighting enhancers using global accessibility score does not need any inter-cell tag-count normalisation. Using the first method also makes it possible to have a uniform transformation of scATAC-seq read-count matrix from different scientific groups without re-calculating tag-counts using the aligned DNA fragments (bam or sam files) on a common peak-list.

The second method can be used while analysing groups of cells with high heterogeneity among each other. In the second method, the quantile-normalisation of read-counts of cells is performed, followed by the division of read-count of every genomic site by its mean read-count across all the cells. The second method needs heterogeneity among cells in order to be more effective.

For every cell, the peaks having high normalised tag-count are selected and used as a set of positives (foreground) and the set of all peaks is used as background. Usually, we use a threshold of 1.25 above global accessibility score for choosing foreground peak, but it could vary depending on stringency needed. The chosen peaks in the positive set are highly likely to be enhancers and regulatory sites with cell-type-specific activity. Then for every peak, a most proximal gene within 1Mbp is found, and peaks which do not have any gene within 1Mbp is dropped. To decide the most effective statistical test we used two different ways to calculate the significance of enrichment of pathways and benchmarked them by applying them using a known set of markers (gene-sets) for different cell types. The statistical methods we use are binomial and hypergeometric tests. With a binomial test, to calculate statistical significance (P-value) for a gene-set m whose genes appear proximal to k_m_ out of n peaks in foreground set, we use the formula below

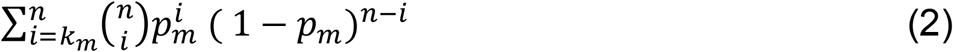

Here p_m_ represents the probability of genes from the gene-set m to appear as proximal to peaks in the background list. With the hypergeometric test, the calculation of statistical significance (P-value) is done using the formula

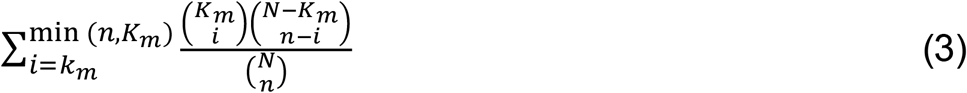

Where K_m_ is the number of times genes of gene-set m appear as proximal to peaks in the background, and N is the total number of peaks in the background set. As above k_m_ represents the number of times out of n foreground peaks, the proximal genes are from gene-set m.

### Normalisation free Gene-set enrichment for single-cell expression

For estimating the significance of enrichment of pathway (gene-set) using scRNA-seq, we use the logarithmic value of the expression (FPKM, TPM, RPKM, UMI-count) of genes and treat every cell independently from each other. Thus unlike other published methods, we avoid creating artefacts which can happen due to the unresolved issue of estimating the distribution of tag-count of a gene across multiple samples (or cells) for normalisation. As scaling and normalisation across different cells can create artefacts due to a variable level of noise and gene dropout rate among them. Rather, we use the widely accepted fact that within a sample (cell) non-zero expression (FPKM, TPM and RPKM) values of genes follow approximately log-normal distribution (see Figure S1). We treated UMI-counts as expression, as UMI-counts are independent of gene-length bias[12]. We modelled the distribution of log(gene-expression) as bimodal such that one mode corresponds to genes with zero count and other mode correspond to a normal distribution for genes with non-zero expression. Thus, probability distribution function (pdf) for log(expression) value x in a cell can be written as

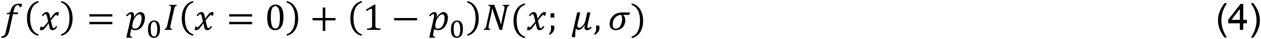

Where N(x; μ, σ) represent Gaussian pdf for genes with non-zero expression (FPKM, TPM, UMI-count) and I(x = 0) is the indicator function, whereas p_0_ represents a fraction of genes with zero expression-value. Here, the fraction of genes with zero expression-value represents both true low expression as well as dropout (undetected expression). The variables μ and σ represent the mean and standard deviation respectively of logarithmic values of only non-zero expression −(FPKM, TPM, UMI-count) in a single-cell. Thus, for every cell, we use its own value of μ and σ to convert the log scale value of non-zero expression of a gene into P-value (right-tailed) assuming Gaussian distribution. The reason for the conversion of gene-expression value to P-value is that it gives an advantage of combining them using Brown’s method that can be used for P-values derived using Gaussian distribution. Thus we combine p-values of genes belonging to a gene-set using Brown’s method[55]. Brown’s method is meant to combine p-values which have a dependence upon each other. Using Brown’s method the combined p-value for a gene-set with k genes with non-zero expression can be given by

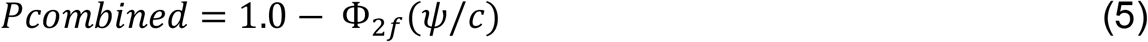

Where 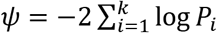 such that *P*_*i*_ is the p-value of log(expression) of gene i in a sample/cell and Φ_2*f*_ is the cumulative distribution function for the chi-square distribution 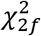. Here *f* is the scaled degree of distribution and is calculated as *f* = *E*[*ψ*]^2^/*var*[*ψ*] [55]. The value of *c* in equation (5) is calculated as

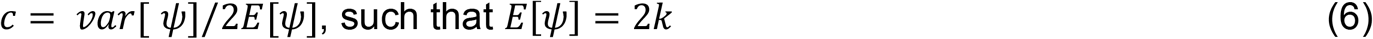

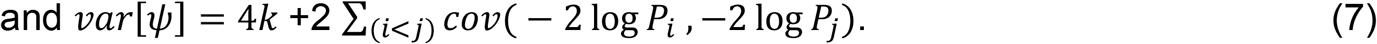

This procedure leads to the calculation of combined P-value for each gene-set in every cell. Covariance between log-p-values of genes is calculated by using their values in all the cells in the same data-sets. In order to have robust estimate not affected by just 1 or 2 genes, we use a threshold of minimum 5 genes with non-zero expression to calculate a combined p-value for a gene-set. However, combined p-values could also have many unwanted effects from house-keeping genes, promiscuously enriched gene-set and multiple hypothesis testing. Hence, we corrected the p-values with a permutation-based test using a null model.

In order to make a null model, we first randomly chose cells from multiple studies so that we can have an equal representation of multiple cell-type. The scRNA-seq profiles of these studies (table S5) were downloaded from recount2 database[56], which provides counts/expression on the same ensemble gene list. The ensemble gene ids were converted to official gene symbols. Those ensemble genes whose official symbol was not available were dropped from the list. We performed hierarchical clustering of cells using genes selected on the basis of the criterion of coefficient-of-variation [57]. We choose highly variable genes based upon the coefficient of variation in each bin when the genes are binned according to their mean value across all cells. In other words, we chose 500 features only for hierarchical clustering while making null model. Using dynamic cutting of the hierarchical tree, we achieved clusters (or classes) of cells. We chose 1000 pairs of cells such that in a pair the cells belonged to different clusters in hierarchical clustering based result. It was done to ensure heterogeneity in the null model. For each pair, we took an average expression value for all gene. Thus, the null model consisted of 1000 expression vectors (false cells), each being average of gene-expression profiles of two cells. For every false-cell vector in the null model, the combined p-values of gene-sets were calculated using the method mentioned above. Thus, for every pathway (gene-set) we achieved 1000 p-values corresponding to the number of false-cells in the null model. For a pathway (gene-set) to calculate adjusted P-value in the target cell, we take the proportion of cells in the null model, which had lower combined P-value than the target cell.

### UniPath’s approach of temporal ordering of cells using pathway scores

Nearly all the methods developed so far for temporal ordering of single-cells use gene-expression or read-count data. Hence to utilize the continuity in pathway activity among cells and to get better insight using single-cell profiles, we developed a novel temporal ordering method which can work efficiently using the pathway scores of single-cells. Our method first performs hierarchical clustering of cells before finding the order among the clusters of cells, followed by distance weighting and learning minimum spanning tree. As illustrated by Zhicheng and Hongkai, applying minimum spanning tree detection directly on raw distances among cells like monocle-1 [25] can lead to a false connection between cells due to noise or other bias[31]. However, following Zhicheng and Hongkai’s method, it is not feasible to get true ordering at single-cell resolution. Hence, we developed an approach, such that after initial classification of cells using pathway scores, we shrink (or weight) distances among every cell pair based on their belonging-ness to the same class and using neighbourhood index among their classes. To calculate neighbourhood index among classes, we first find top k nearest neighbour for every cell. Then for every class, we count the number of times its cells have top k neighbours in other classes. For example, if cells in class A has total M neighbours in others classes out of which mb cells are from a class B then we calculate neighbourhood index of A with B (A ->B) as mb/M. We shrink the distances between the cells in class A and class B by mb/M. After two stages of shrinkage of distances among cells, we use shrinked-distance matrix to find a minimum spanning tree. We plot the minimum spanning tree using the netbiov R library[58]. The minimum spanning tree drawn using our approach has fewer chances to be influenced by noise as the distances among cells are shrunk using consensus information.

### Test for pathway score accuracy and consistency of UniPath

Even though we tested UniPath and PAGODA using cell-type markers, we also used spike-in method for measuring accuracy for pathway scores. For this, first, we collected gene-sets for non-immune pathways. In this collection, we also added 2 gene-set specific to B cells and 2 pathways for T cells. Both PAGODA and UniPath were used with collected gene-set on FPKM of B and T cells. UniPath and PAGODA were evaluated based on the presence of relevant pathway among top 5 gene-sets in their output.

### Simulating systematic dropout

There could be many reasons for zero read-count for genes in scRNA-seq data-set. Those reasons include true biological cause and random and systematic dropout. Unlike random dropout, systematic dropout often leads to errors during dimension reduction and classification. Therefore, we simulated systematic dropout to evaluate UniPath. To create systematic dropout for evaluation (shown in Fig. 3a) using a combined dataset of hESC and Myoblast cells [24, 25], we first randomly chose some genes (10% genes). We dropped the FPKM values of chosen genes from randomly selected hESCs; in other words, we made the FPKM values of chosen genes to zero in a few hESCs. We varied the percentage of genes whose FPKM could be dropped to test the robustness of UniPath. The case of systematic dropout is actually different from most of the technical batch effect.

### Differential co-occurrence analysis

We use a permutation test to estimate the significance of the difference of pathway co-occurrence among two groups of cells. For a pathway pair, we first calculate the difference between spearman correlation values of their enrichment scores (adjusted P-value) in two groups of cells. We call it a true difference. We perform random-shuffling of group-labels of cells and calculate the difference in spearman-correlation of enrichment scores among two shuffled groups. Thus, for a pair of pathways, we make a collection of a set of false-difference in correlations using shuffled groups. The p-value is calculated as a fraction of false-differences which are greater than the true difference in term of absolute value. Notice that, here, we use spearman correlation of adjusted P-value (pathway score), not just the combined p-value of gene-sets. Using adjusted P-value increases robustness as it becomes rank based scores which helps in filtering out effect due to only 1 or 2 genes. Thus, if two pathways are correlated using their adjusted P-value, the correlation has less chance to be affected by only 1 or 2 genes or outliers. Even though random permutation-based significance test reduces possible covariates, we have ignored differential co-occurrence among gene-sets which have high Jaccard index for overlap of genes. For hESC cells differentiating towards DE, we performed differential co-occurrence analysis for every time point of differentiation. Such as for differential co-occurrence analysis for time point of 12 hours, we compared them with cells not belonging to 12 hours.

### Single-cell expression profiling for non-small lung cancer cells

The source and culture condition for Tumour sphere (TS) and Adherent cells(Adh) are mentioned in Wang et al. [42]. Tumour sphere (TS) line derived from lung cancer patient were maintained in medium with DMEM/F12 (US Biomedical), 4mg/ml Bovine Serum Albumin (Sigma), Non-essential amino acids, sodium pyruvate (Life Technologies) and 20ng/ml Epidermal Growth Factor, 4 ng/ml bovine Fibroblast Growth Factor and Insulin – Transferrin Selenium (Sigma).

Tumour sphere derived adherent (Adh) cells were grown in the same media as above, without EGF, bFGF, ITS and BSA. For Adh cells, media was supplemented with 10% fetal bovine serum.

### RNA extraction, library construction, sequencing for NSCLC cells

NSCLC single-cells in suspension were dissociated using trypsin and loaded into C1 96 well-integrated microfluidic chip (IFC) as per manufactures guidelines. The single-cells were captured in C1 96 (large size) IFC using Fluidigm-C1 system. The captured single-cells were imaged using auto imaging fluorescent microscope to identify the viable single-cells and to omit the doublets. The reverse transcription and cDNA pre-amplification reagents were prepared using SMART-seq2 protocol and loaded into the IFC. Later reverse transcription and cDNA amplification were processed using SMART-seq2 script automatically in C1-Fluidigm machine. After harvesting cDNA from C1 chip, the samples were quantified using picogreen assay and normalized to the range 0.2-0.3ng/μL. The quality of the cDNA product was verified using high sensitivity DNA assay in Agilent bio-analyzer machine. The harvested single-cell cDNA was barcoded in 96 well plate using Nextera XT Library Prep kit (Illumina). Uniquely barcoded libraries from single-cells pooled together and sequenced using a HiSeq-Hi-output-2500 sequencer (Illumina). In total, there were 87 TS cells and 75 Adh cells.

## Implementation and data availability

UniPath is implemented using R and is available at https://reggenlab.github.io/UniPathWeb/ As well as at https://github.com/reggenlab/UniPath

The FPKM data for single-cell RNA-seq for lung cancer cells is available with UniPath package. The raw sequences are being uploaded to BioSample database. The raw sequences contain genomic identity information of cancer patients; hence they can be accessed only with permission.

## Authors contributions

VK and DS designed the study, WLT helped in designing study related to cancer cells. VK and SC wrote the code and the manuscript. SC and VK also executed the code on data-sets for generating results and figures. SS and SLK prepared the library for single-cell RNA-seq of lung-cancer cells. ZW and WLT cultured non-small cell lung cancer cell-line and provided information about cultured cells and their behaviour.

## Acknowledgement

This project was supported by YIG Grant (BMRC/YIG/1510851023) provided to Vibhor Kumar by BMRC, A-STAR, Singapore.

## Ethics declarations

we used a previously characterized LC32 cell lines derived from resected primary non-small-cell lung cancer (NSCLC) adenocarcinoma samples

