## supplementary methods, table for "UniPath: A uniform approach for pathway and gene-set based analysis of heterogeneity in single-cell epigenome and transcriptome profiles"

Figure S1

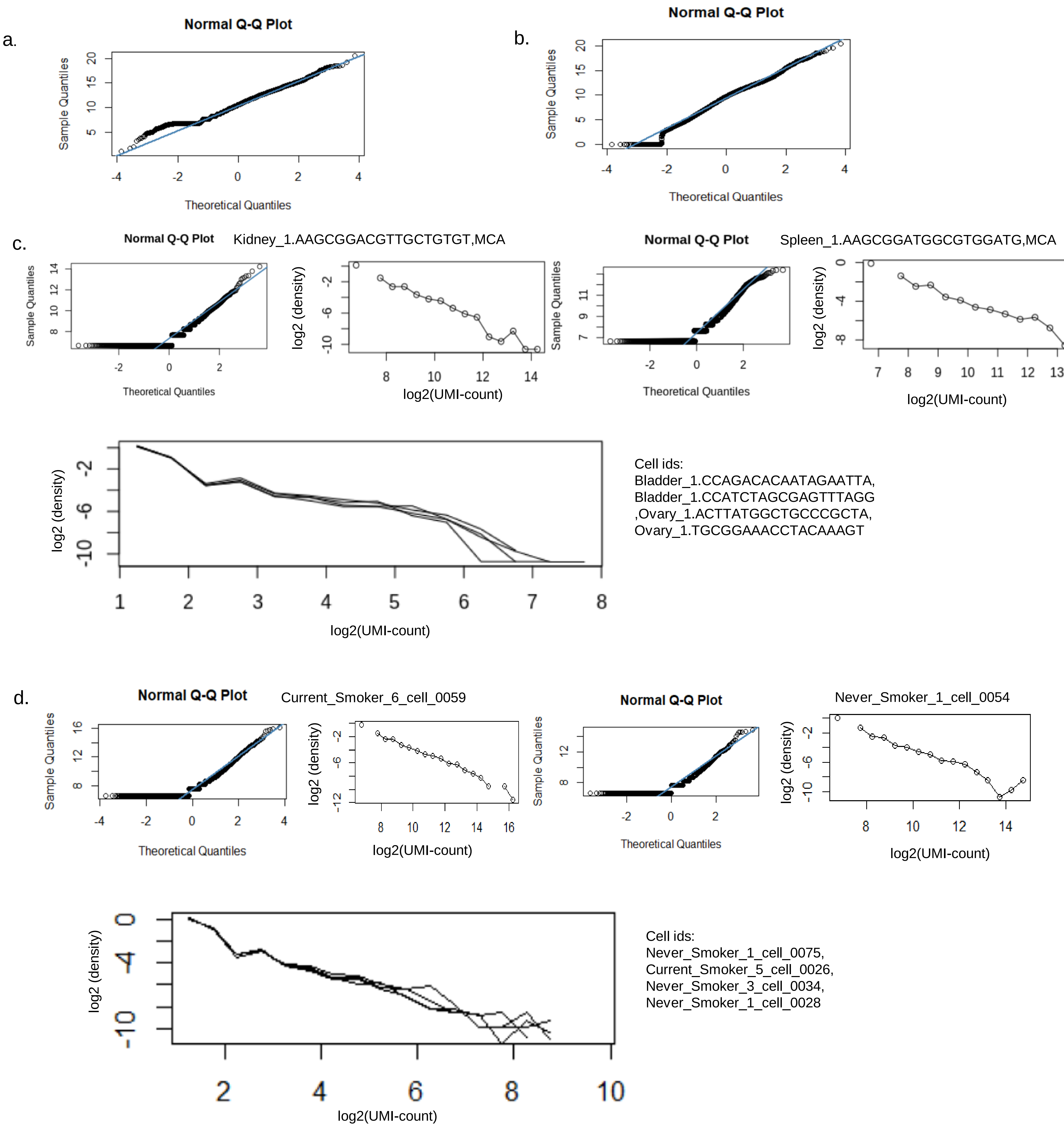


**Figure S1: Towards fitting a distribution for non-zero gene-expression values in a cell. (a)** Quantile-quantile (q-q) plot of log2(TPM) of genes with non-zero expression in a cell with single cell RNA seq profile (GSE75748). **(b)** q-q plot for log2 (FPKM) of genes in a cell using its single cell RNA seq profile (GSE52529). **(c)** q-q plot of log2(UMI-count) for genes with non zero UMI in single cell RNA-seq profile from mouse cell atlas data-set (GSE108097). The log2(density) is also plotted w.r.t log(UMI-count) on x axis. The plot shows almost a straight line, supporting the assumption of log-normal distribution for UMI-count. **(d)** q-q plot and log2 density plot for non zero UMIs using single cell RNA seq data (GSE131391). Overall q-q plot and log2(density) plot for UMI-count from two different data-set hints it approximately fit log-normal distribution.

Figure S2

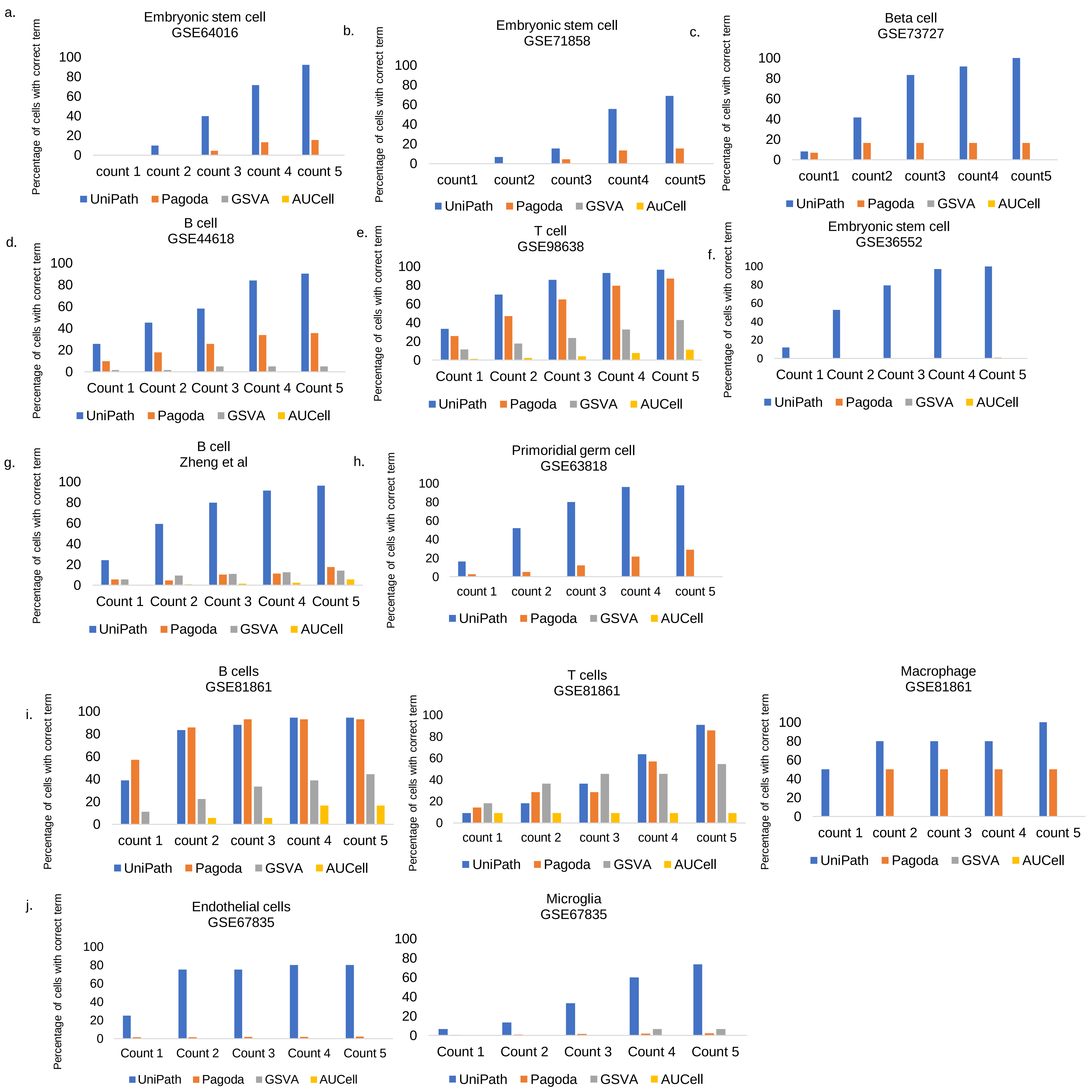


**Figure S2**: Comprehensive evaluation of UniPath for single-cell RNAseq for highlighting correct terms in top enriched results. The terms here are cell-types and gene-sets of terms are set of marker genes for corresponding cell-type.**(a)** Embryonic stem cell (ESC) percentage detected in homogeneous dataset of hESC scRNAseq (Fluidigm C1 platform, non-UMI). The bars show percentage of cells with correct cell-type among top enriched terms. 'count1' shows percent of cells with correct cell-type as the first enriched term. Similarly count5 shows percentage of cells with correct cell-type among top-5 enriched terms. **(b)** Accuracy for cell-type detection in homogeneous dataset of ESC processed using FRISCR and TritonX-100 Lysis. (Using TPM, non-UMI) **(c)** correct cell-type detection percentage for heterogeneous dataset of Beta Cell (Using RPKM, non-UMI) **(d)** correct cell-type detection percentage for homogeneous dataset of B-cell (Smart-seq protocol, non-UMI) **(e)** percentage of correct detection in homogeneous scRNAseq dataset of T-cell (Smartseq2 and Tang et al., 2010 protocol). **(f)**. For heterogeneous dataset of Embryonic stem cell (Tang protocol, non-UMI). **(g)** correct detection in homogeneous dataset of B-cell (non-UMI) **(h)** Primordial germ cell ; heterogeneous dataset (Tang protocol, non-UMI). **(i)** Cell type detection for heterogeneous dataset of B-cell, T-cell and macrophages (fluidigm based scRNA-seq protocol, non-UMI). **(j)** Detection of cell-types from scRNAseq profiles of Endothelial and Microglial cell processed using fluidigm C1 based protocol (non-UMI).

### Figure S3

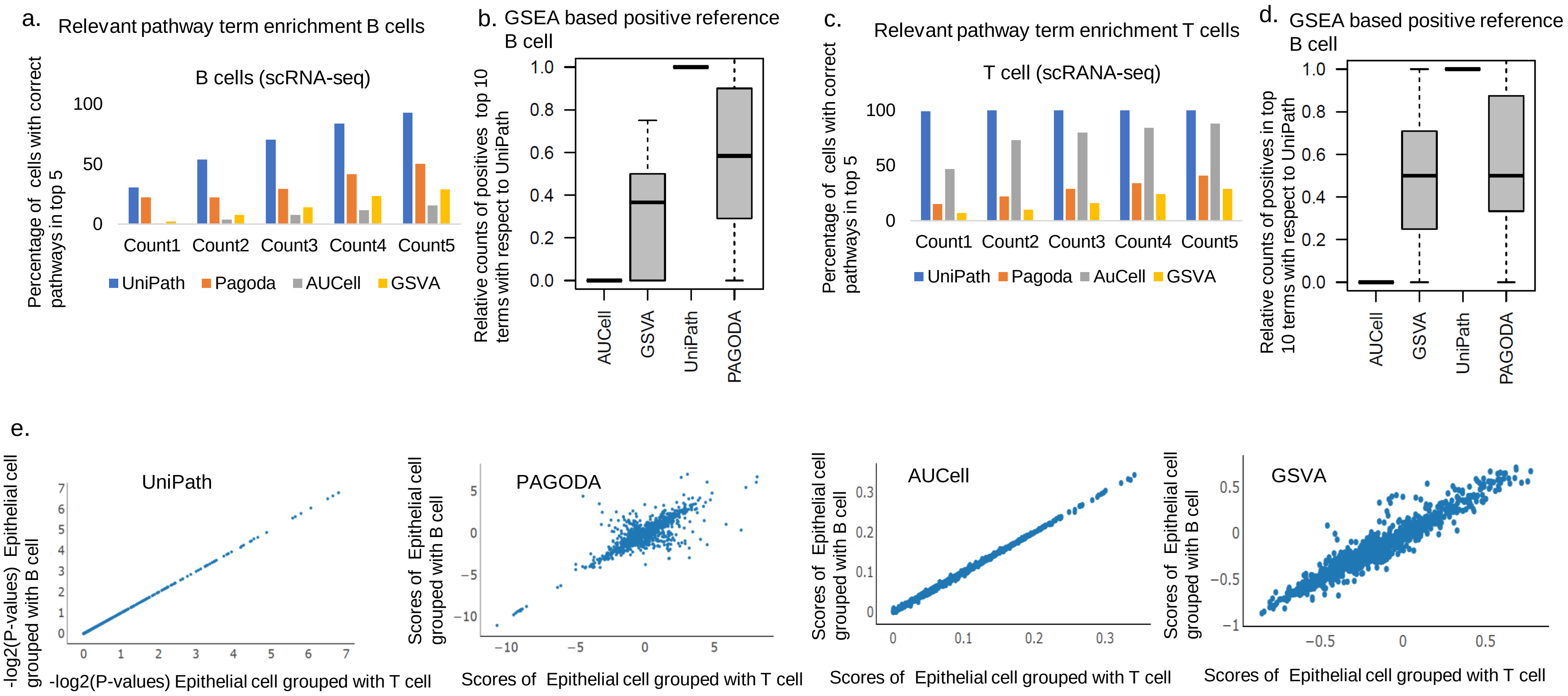


**Figure S3: Evaluation of** consistency and enrichment calculation for pathways by UniPath ,PAGODA, AUCell and GSVA. **(a) Evaluation of e**nrichment of B cell pathways where 'Count1' shows percentage of cells with correct first enriched term. Here the list of gene-set consisted of non-immune pathways and two pathways (as positives) known to be active in B cell. (b) Boxplots shows relative counts of positives in top 10 terms based on scores calculated by four different methods. Here the list included all gene-sets (terms). The positives are those gene-sets which were found to be enriched in B cell group w.r.t other cell types in mouse cell atlas data-set by GSEA. The counts of positives in each cell was normalized by counts (of positives) found using UniPath in same cell. **(c)** Enrichment of T cell pathways where 'Count1' shows percentage of cells with correct first enriched term. (d) Boxplots showing relative counts of positives in top 10 terms. The positives are those gene-sets which were found to be enriched in T cell group in mouse cell atlas data-set by GSEA. The count of positives in each cell is normalised by corresponding values for UniPath based results in same cell. **(a)** Scores of Epithelial cells calculated by UniPath, PAGODA, AUCell and GSVA when grouped with B and T cells respectively. The data-set used here was adapted from study with GEO ID: GSE81861. Output of UniPath and AUCell for a cell is consistent and is not affected by the type of neighboring cells. Whereas for PAGODA the estimate of dispersion (equivalent to enrichment) for pathway is dependent upon composition of cell-type in the data-set.

Figure S4

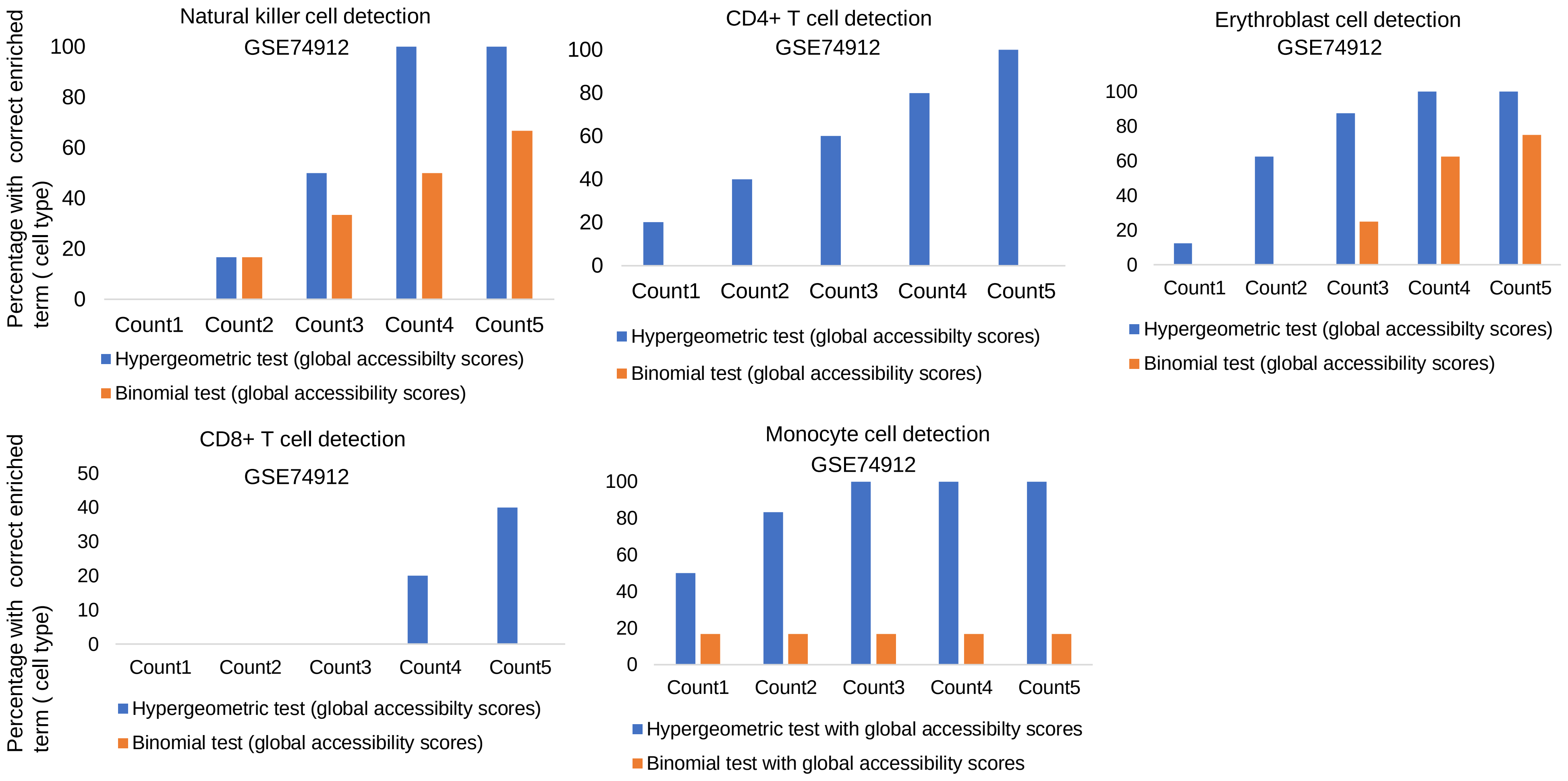


**Figure S4:** Evaluation of accuracy of UniPath for estimating gene-set enrichment using open-chromatin profile of bulk sample of cell lines. The gene-sets of marker for cell-types are used here for benchmarking. Shown here is percentage of cells with correct cell-type gene-set among top enriched terms. ‘count1’ shows percent of cells with correct cell-type as the first enriched term. Similarly, count3 shows percentage of cells with correct cell-type among top-3 enriched terms. The bulk ATAC-seq were adapted from study with (GEO ID: GSE74912). Enrichment score for Natural killer cell, CD4+ T cell, CD8+ T cell, Monocyte and Erythroblast cells were calculated by UniPath using hypergeometric and binomial test. For every cell-type results are shown for both situation, when enhancers were enriched using division global accessibility scores.

Figure S5

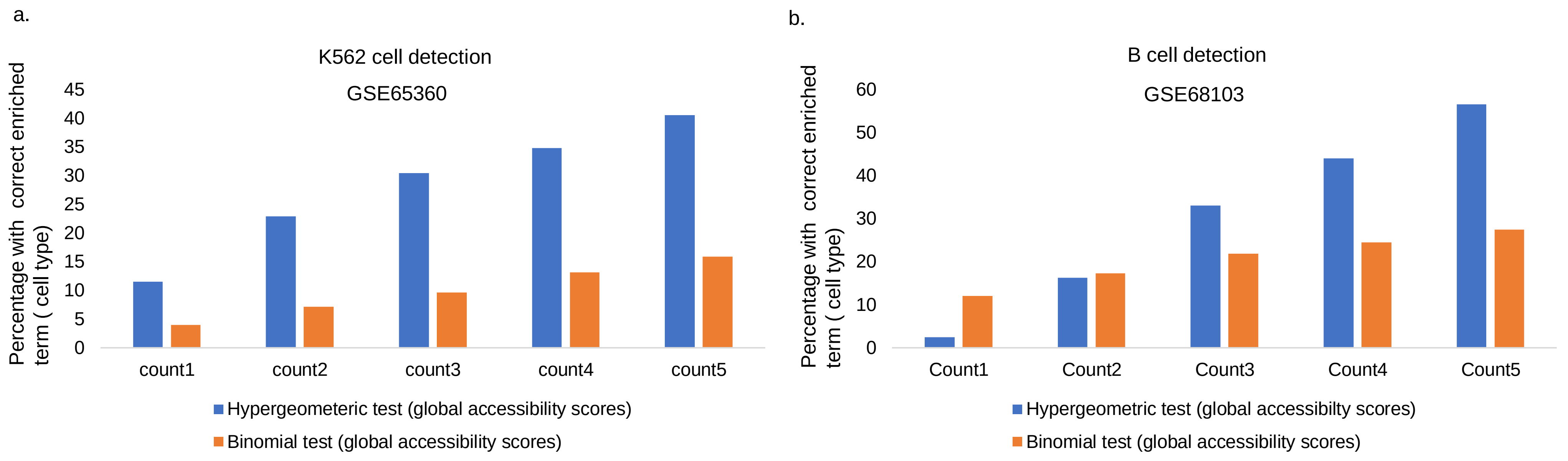


**Figure S5:** Accuracy of UniPath for enrichment of correct gene-set in top results for single cell open-chromatin profile. Evaluation of accuracy is done using marker gene-set for cell-types while applying UniPath on scATAC-seq profiles. **(a)** K562 cell detection using hypergeometric test and binomial test using enhancer highlighted using global accessibility scores ( data set GEO ID: GSE65360). **(b)** percentage of Correct cell-type detection using scATAC-seq of B-cell (GEO ID: GSE68103).

### Figure S6

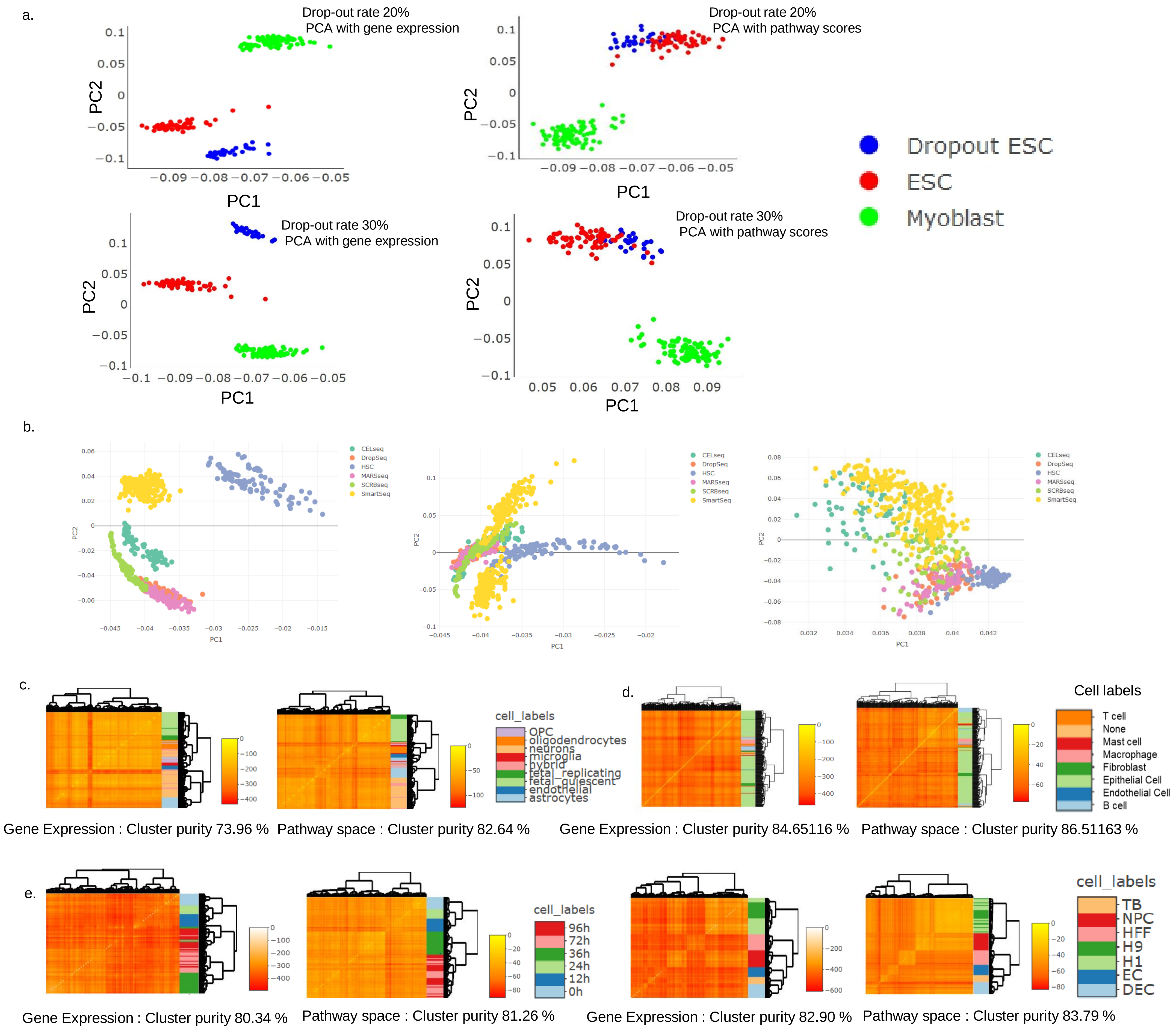


**Figure S6**: Batch correction by UniPath and clustering purity using pathway scores **(a)** Combined human ESCs from a study (GSE75748) and Myoblast from another study (GSE52529). Few ESCs were subjected to systematic drop-out of 10% and 30%. Principal component analysis (PCA) based dimension reduction and visualization using gene expression shows two separate clusters of human ESC. However UniPath is robust to systematic variability in drop-out rate of 20% and 30%. In PCA based dimension-reduction of pathway scores by UniPath, all the hESC cells come together in one group. **(b)** PCA based visualization of batch effect in mouse embryonic stem cell lines processed using various protocols (GSE75790) combined with Hematopoietic stem cells from study GSE71794 in gene expression space. Batch effect removal in gene expression space using Limma. Batch effect removal done in pathway space using Limma shows mixing of ESCs to some extent and separation of HSCs. **(c)** Heatmaps showing comparable clustering purity in gene expression and pathway space for various cells types from study GSE67835. **(d)** Heatmaps showing comparable clustering purity in gene expression and pathway space for various cells types from study GSE81861. **(e)** Heatmaps showing comparable clustering purity in gene expression and pathway space for various cells types from study GSE75748.

Figure S7

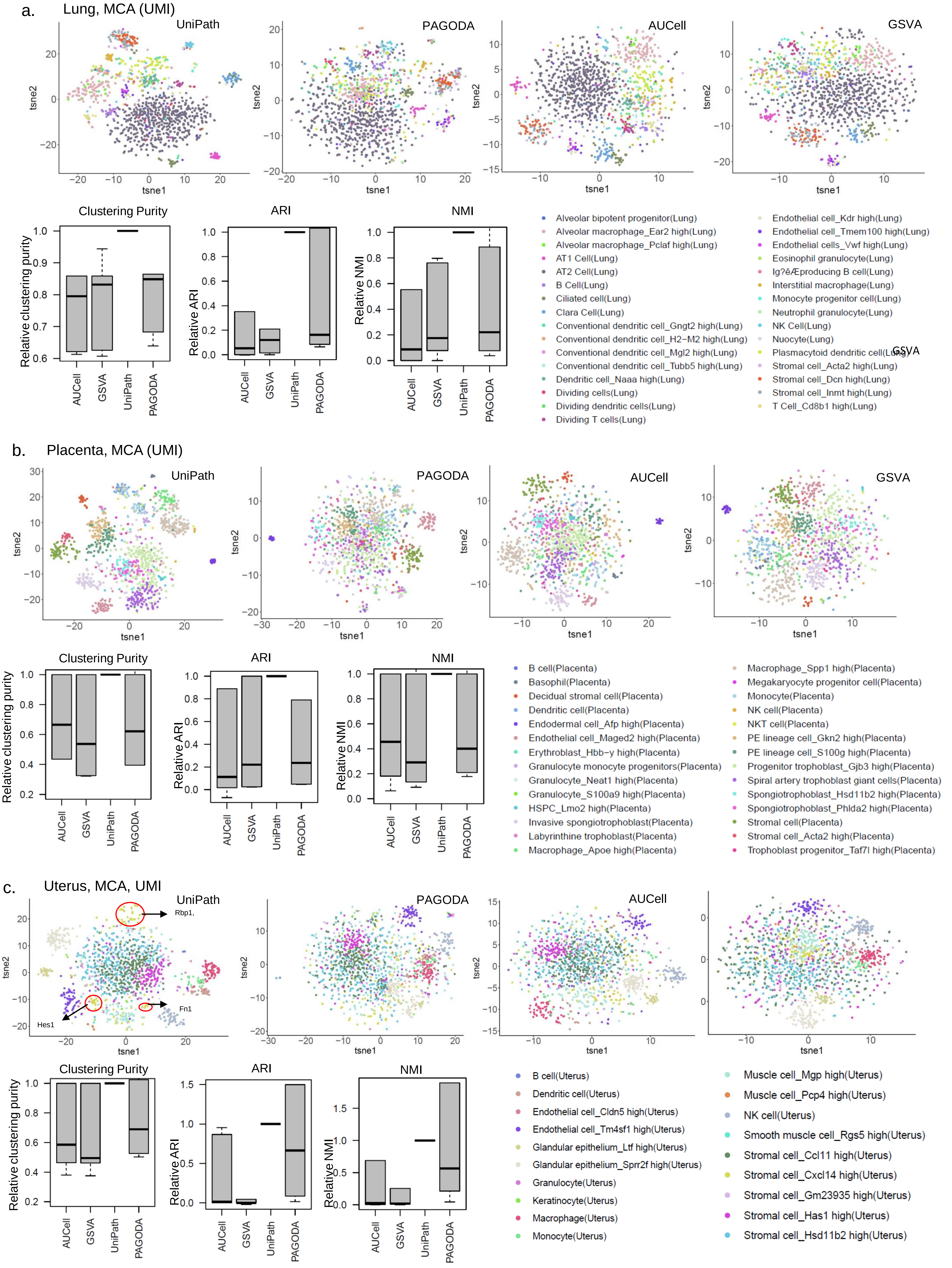

a.
Lung, MCA (UMI)
UniPath
PAGODA
AUCell
GSVA
tsne1
tsne2
Clustering Purity
ARI
NMI
Relative clustering purity
Relative ARI
Relative NMI
AUCell
GSVA
UniPath
PAGODA
Alveolar bipotent progenitor(Lung)
Alveolar macrophage_Ear2 high(Lung)
Alveolar macrophage_Pclaf high(Lung)
AT1 Cell(Lung)
AT2 Cell(Lung)
B Cell(Lung)
Ciliated cell(Lung)
Clara Cell(Lung)
Conventional dendritic cell_Gngt2 high(Lung)
Conventional dendritic cell_H2−M2 high(Lung)
Conventional dendritic cell_Mgl2 high(Lung)
Conventional dendritic cell_Tubb5 high(Lung)
Dendritic cell_Naaa high(Lung)
Dividing cells(Lung)
Dividing dendritic cells(Lung)
Dividing T cells(Lung)
Endothelial cell_Kdr high(Lung)
Endothelial cell_Tmem100 high(Lung)
Endothelial cells_Vwf high(Lung)
Eosinophil granulocyte(Lung)
Ig?êÆproducing B cell(Lung)
Interstitial macrophage(Lung)
Monocyte progenitor cell(Lung)
Neutrophil granulocyte(Lung)
NK Cell(Lung)
Nuocyte(Lung)
Plasmacytoid dendritic cell(Lung)
GSVA
Stromal cell_Acta2 high(Lung)
Stromal cell_Dcn high(Lung)
Stromal cell_Inmt high(Lung)
T Cell_Cd8b1 high(Lung)
b.
Placenta, MCA (UMI)
UniPath
PAGODA
AUCell
GSVA
tsne1
tsne2
Clustering Purity
ARI
NMI
Relative clustering purity
Relative ARI
Relative NMI
AUCell
GSVA
UniPath
PAGODA
B cell(Placenta)
Basophil(Placenta)
Decidual stromal cell(Placenta)
Dendritic cell(Placenta)
Endodermal cell_Afp high(Placenta)
Endothelial cell_Maged2 high(Placenta)
Erythroblast_Hbb−y high(Placenta)
Granulocyte monocyte progenitors(Placenta)
Granulocyte_Neat1 high(Placenta)
Granulocyte_S100a9 high(Placenta)
HSPC_Lmo2 high(Placenta)
Invasive spongiotrophoblast(Placenta)
Labyrinthine trophoblast(Placenta)
Macrophage_Apoe high(Placenta)
Macrophage_Spp1 high(Placenta)
Megakaryocyte progenitor cell(Placenta)
Monocyte(Placenta)
NK cell(Placenta)
NKT cell(Placenta)
PE lineage cell_Gkn2 high(Placenta)
PE lineage cell_S100g high(Placenta)
Progenitor trophoblast_Gjb3 high(Placenta)
Spiral artery trophoblast giant cells(Placenta)
Spongiotrophoblast_Hsd11b2 high(Placenta)
Spongiotrophoblast_Phlda2 high(Placenta)
Stromal cell(Placenta)
Stromal cell_Acta2 high(Placenta)
Trophoblast progenitor_Taf7l high(Placenta)
c.
Uterus, MCA, UMI
UniPath
Rbp1,
Hes1
Fn1
PAGODA
AUCell
tsne1
tsne2
Clustering Purity
ARI
NMI
Relative clustering purity
Relative ARI
Relative NMI
AUCell
GSVA
UniPath
PAGODA
B cell(Uterus)
Dendritic cell(Uterus)
Endothelial cell_Cldn5 high(Uterus)
Endothelial cell_Tm4sf1 high(Uterus)
Glandular epithelium_Ltf high(Uterus)
Glandular epithelium_Sprr2f high(Uterus)
Granulocyte(Uterus)
Keratinocyte(Uterus)
Macrophage(Uterus)
Monocyte(Uterus)
Muscle cell_Mgp high(Uterus)
Muscle cell_Pcp4 high(Uterus)
NK cell(Uterus)
Smooth muscle cell_Rgs5 high(Uterus)
Stromal cell_Ccl11 high(Uterus)
Stromal cell_Cxcl14 high(Uterus)
Stromal cell_Gm23935 high(Uterus)
Stromal cell_Has1 high(Uterus)
Stromal cell_Hsd11b2 high(Uterus)

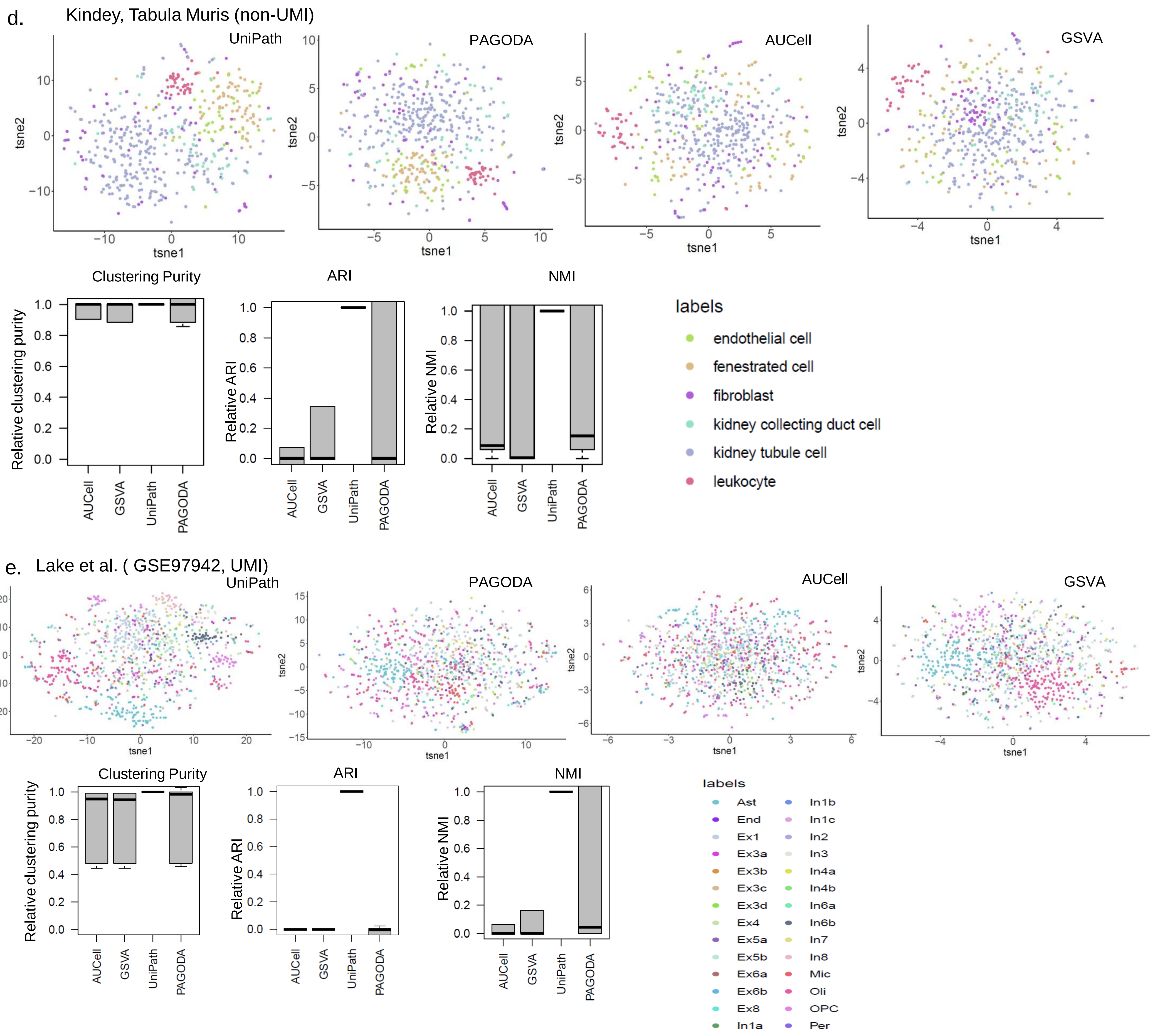


**Figure S7: Quantification of clustering purity and low dimension visualization of scores calculated using UniPath, PAGODA, AUCell and GSVA from scRNA-seq data. (a)** t-SNE for cells from lung (MCA dataset, UMI-counts) and boxplot showing relative clustering purity, ARI and NMI with respect to UniPath calculated using dbscan with different eps values. **(b)** t-SNE for placenta cells (MCA dataset, UMI-counts) and boxplot showing relative clustering purity, ARI and NMI with respect to UniPath calculated using dbscanat various eps values. **(c)** t-SNE of Uterus cells (MCA dataset, UMI-counts) and boxplot showing relative clustering purity, ARI and NMI with respect to UniPath calculated using dbscan with different eps values. **(d)** t-SNE of Kidney cells (Tabula Muris Consortium, non-UMI ) and boxplot showing relative clustering purity, ARI and NMI with respect to UniPath calculated using dbscanat various eps values. **(e)** t-SNE of Occipital Lobe, Visual Cortex region from adult human brain tissue (Lake et al, GSE97942, UMI-counts) and boxplot showing relative clustering purity, ARI and NMI with respect to UniPath calculated using dbscan with different eps values. The purity of clustering using UniPath based score was better than other 3 tools based (PAGODA, GSVA, AUCell) results.

### Figure S8

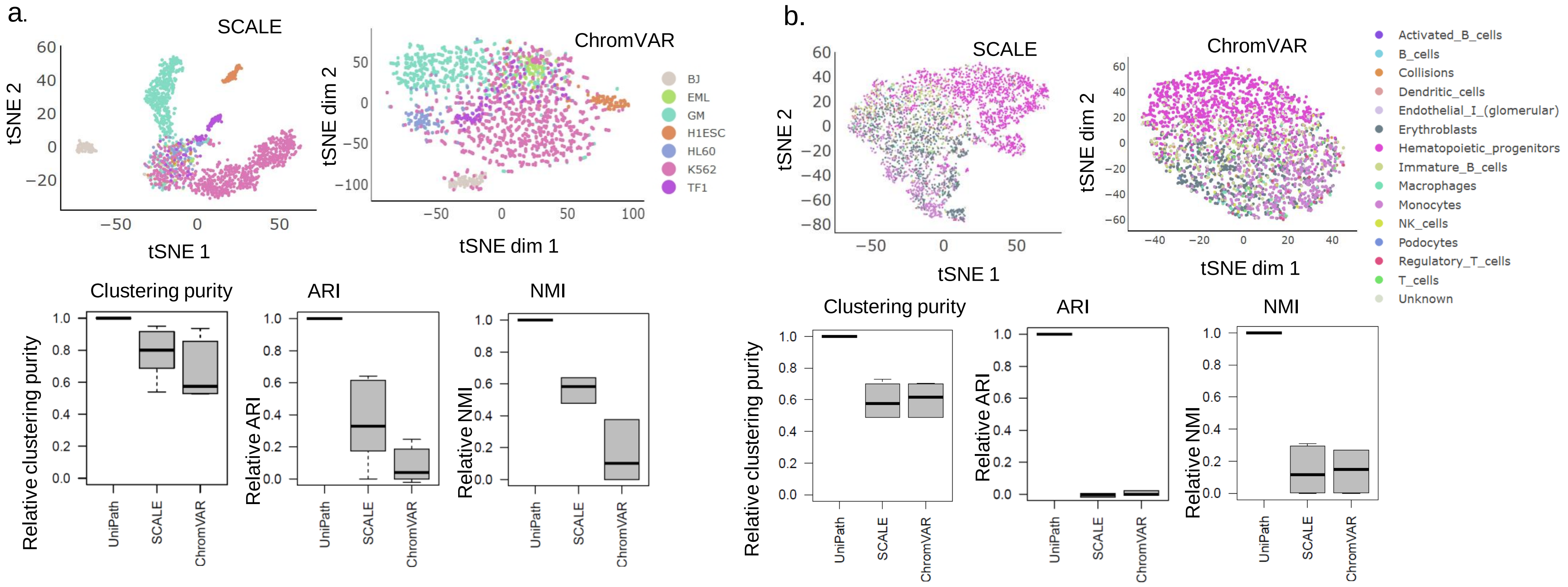


**Figure S8: Quantification of purity of clustering and low dimension visualization of scores calculated using SCALE and chromVar from scATAC-seq data. (a)** t-SNE based scatter plot for scATAC-seq dataset from Buenrostro et al. and boxplot showing relative clustering purity, ARI and NMI with respect to UniPath calculated using dbscan with different eps values. **(b)** Plot of t-SNE coordinates from scATAC-seq dataset from Cusanovish et al. and boxplot showing relative clustering purity, ARI and NMI with respect to UniPath calculated using dbscan with different eps values

Figure S9

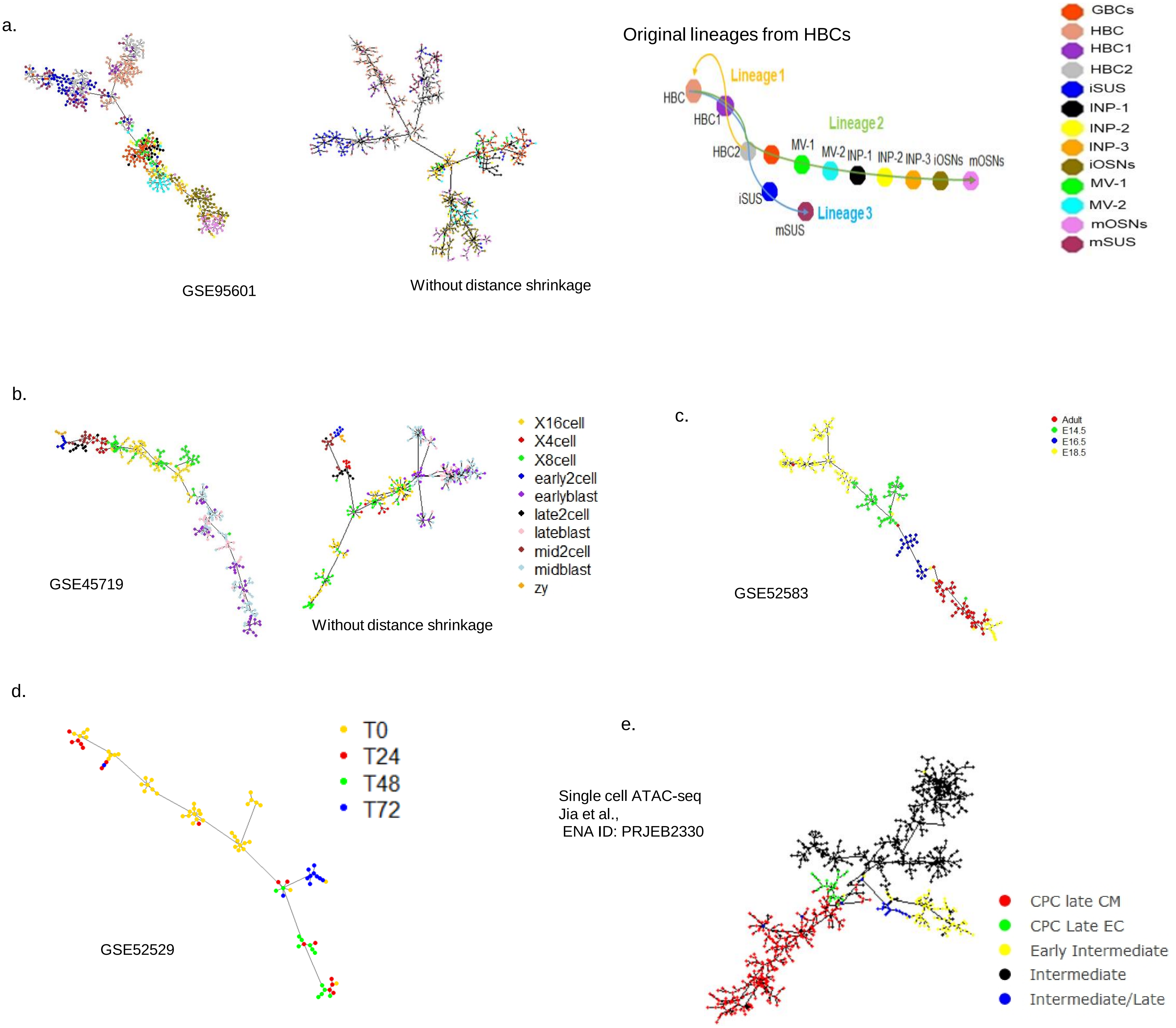


**Figure S9:** UniPath's results for pseudo-temporal ordering of cell using their pathway score. **(a)** Pseudo-temporal ordering for tracing differentiation trajectory of Horizontal basal cell (HBC) to neuronal and sustentacular cell lineages (GEO ID: GSE95601) with and without distance shrinkage. For reference the ideal lineage tree for same data-set has also been shown. **(b)** Pseudo-temporal ordering using pathway scores derived from scRNA-seq profiles of early developmental cells starting from zygote to late blastocyst ( data from Deng et al. data , GEO ID: GSE45719) with and without distance shrinkage. **(c)** Pseudo-temporal ordering showing mouse lung developmental stages (GEO ID: GSE52583). **(d)** Pseudo-temporal ordering showing myoblast cell differentiation at different time points (Trapnell et al, 2014, GEO ID: GSE52529). **(e)** Pseudo temporal ordering in pathway score derived from scATAC-seq profile of cardiac progenitor data (Jia et al., ENA ID: PRJEB23303).

### Figure S10

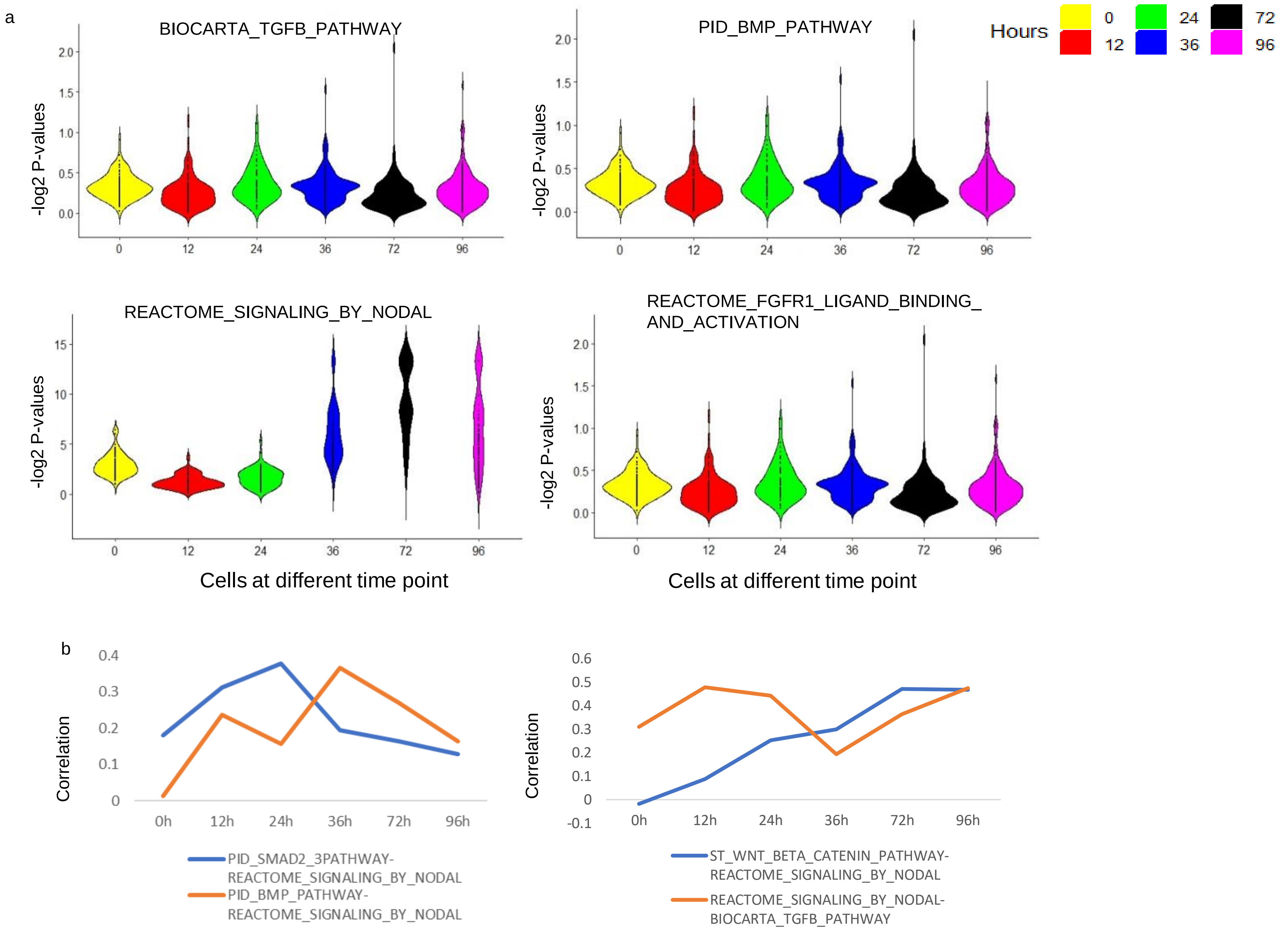


Figure S10. Enrichment and co-occurence of pathways during differentiation towards endoderm (a) Violin plot of enrichment score of pathways at different time point of differentiation. As expected, NODAL signaling pathways had higher score at 72 and 96 hours compared to 0 and 12 hours. Whereas BMP, TGF-beta and FGFR1 signaling pathway scores had bimodal distribution at 36 and 72 hours indicating the heterogeneity in regulation at same time point. (b) Spearman correlation between score of Nodal signalling and other pathways at different time points. Nodal signaling is known to act via smad2/smad3 for differentiation towards mesendodermal lineage (Fei et al. 2010). Here Nodal and smad2 signalling have highest correlation at 24 hours, then it decreased at the start of 36 hours which is consistent with existing literature (Fei, et al, 2010). Similarly Nodal and BMP has highest correlation at 36 hours after which it decreases. BMP is known to support differentiation towards mesendodermal lineage. Decrease in cooccurrence between BMP and nodal signaling at 72 and 90 hours is according the finding reported by Kyle et al. (2013) that high level of BMP inhibits differentiation of primitive streak cells towards definitive endoderm. Correlation between Wnt and Nodal signalling pathway scores increased as differentiation proceeded towards definitive endoderm till 96 hours and such trend support previous reports (Chabra, et al, 2018).

Figure S11

a.

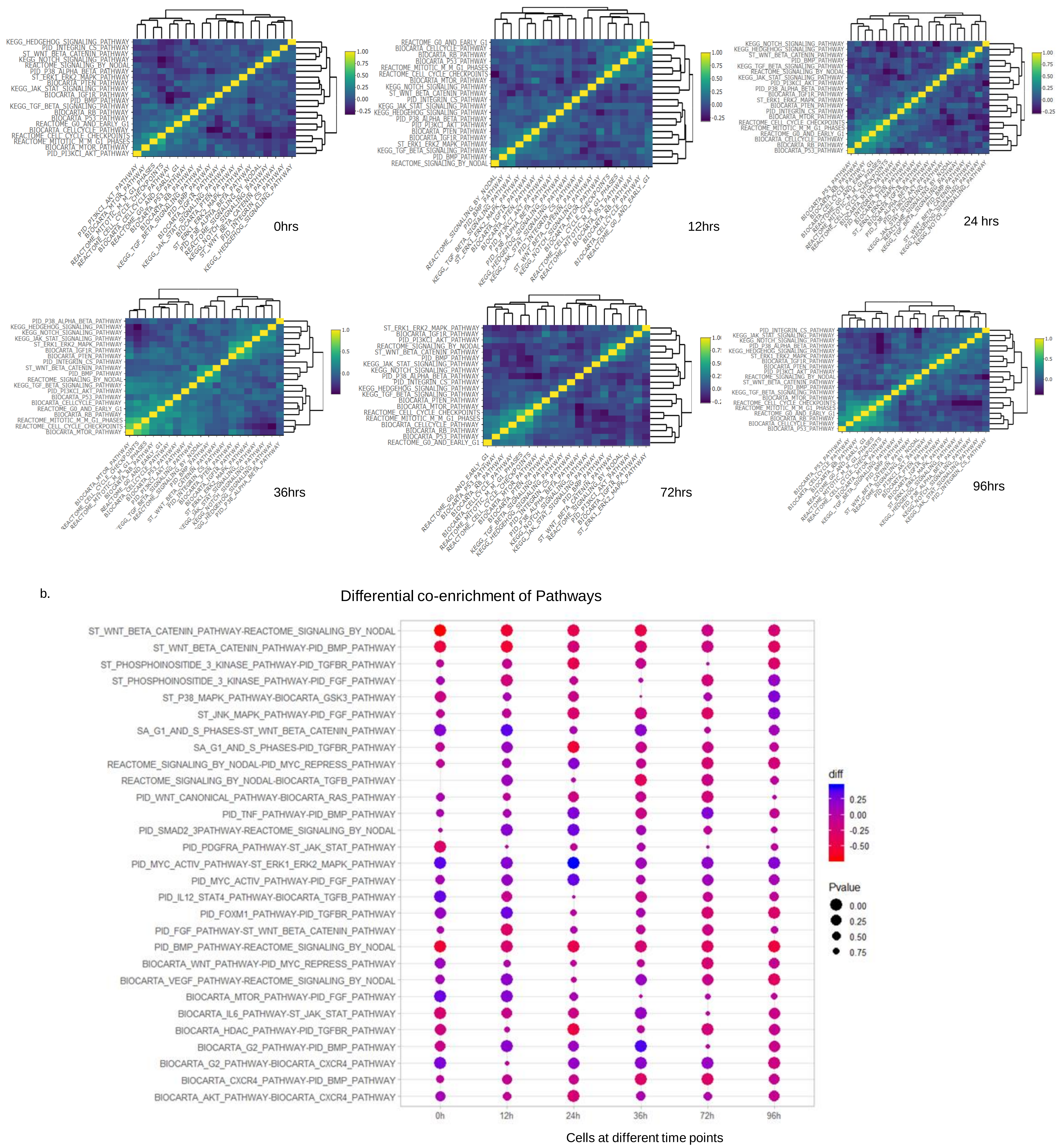


**Figure S11**: Analysis of change in pathway modules and co-occurences in differentiating human embryonic stem cells data (Chu et al, GEO ID: GSE75748) at different time points (0 hours,12 hours, 24 hours, 36 hours and 72 hours. **(a)** Heatmap of correlation between pathways at 6 time of differentiation for differentiating hESC. It shows how modules of pathways change at different time point of differentiation. It can be noticed that Nodal signaling pathways did not co-occur with TGF-beta and BMP pathways at 0 hour. However at 12 hr, 24 hr and 36 hr Nodal, TGF-beta and BMP pathways co-occurred together. **(b)** Differential correlation (co-occurrence/co-enrichment) of pathways, top pathways are selected from each time point and represented by dot-plot. Color of dots represents difference among pathways and size of dot represent P-values. Some of the co-occurrence patterns have been reported previously. Such as FGF induces activation of mTOR and mTOR is involved in involved ESCs pluripotency and suppression of mesoderm and endodermal related activities (Zhou et al., 2009). It can be seen that differential co-occurence of FGF and mTOR pathway is significant at 0 and 12 hours.

### Figure S12

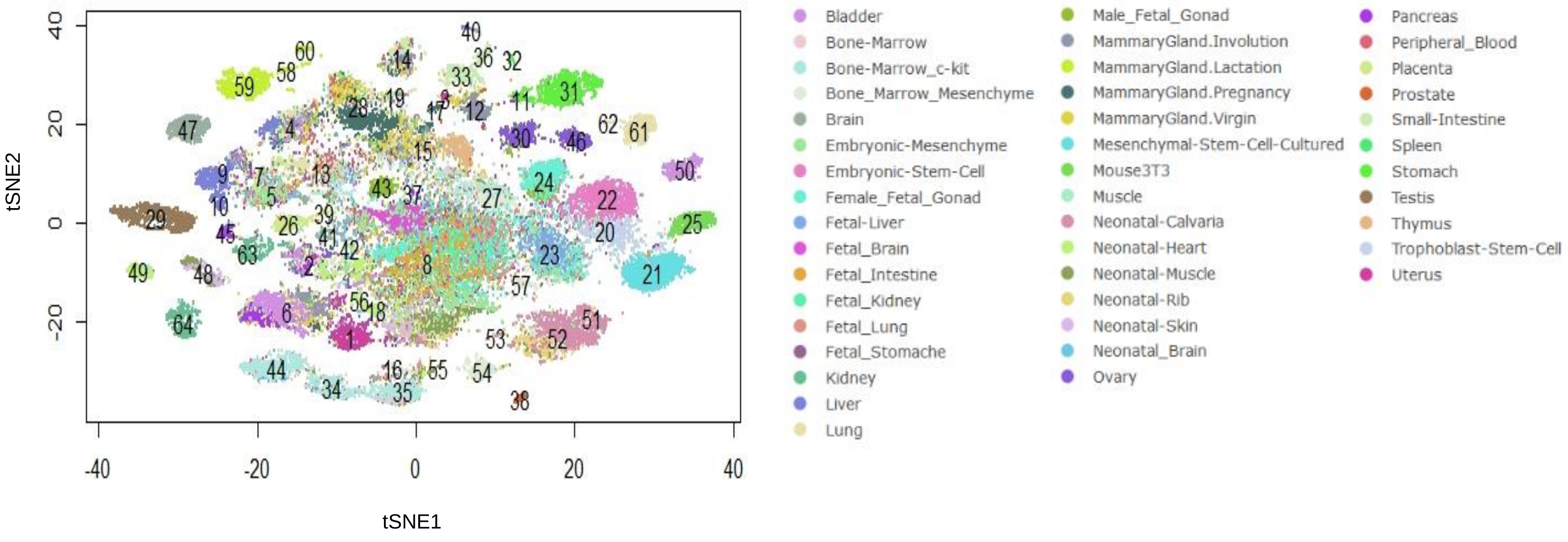


**Figure S12: Pathway scores can be used successfully to distinguish each cell-type as distinct group.** Scatter plot of t-SNE based dimension reduction using pathway scores of single-cells from different organs from mouse cell atlas (Han et al., GEO ID: GSE108097). Scatter plot of t-SNE based dimension reduction of pathway score profile of mouse cell atlas (MCA). The major cluster numbers are also shown

Figure S13

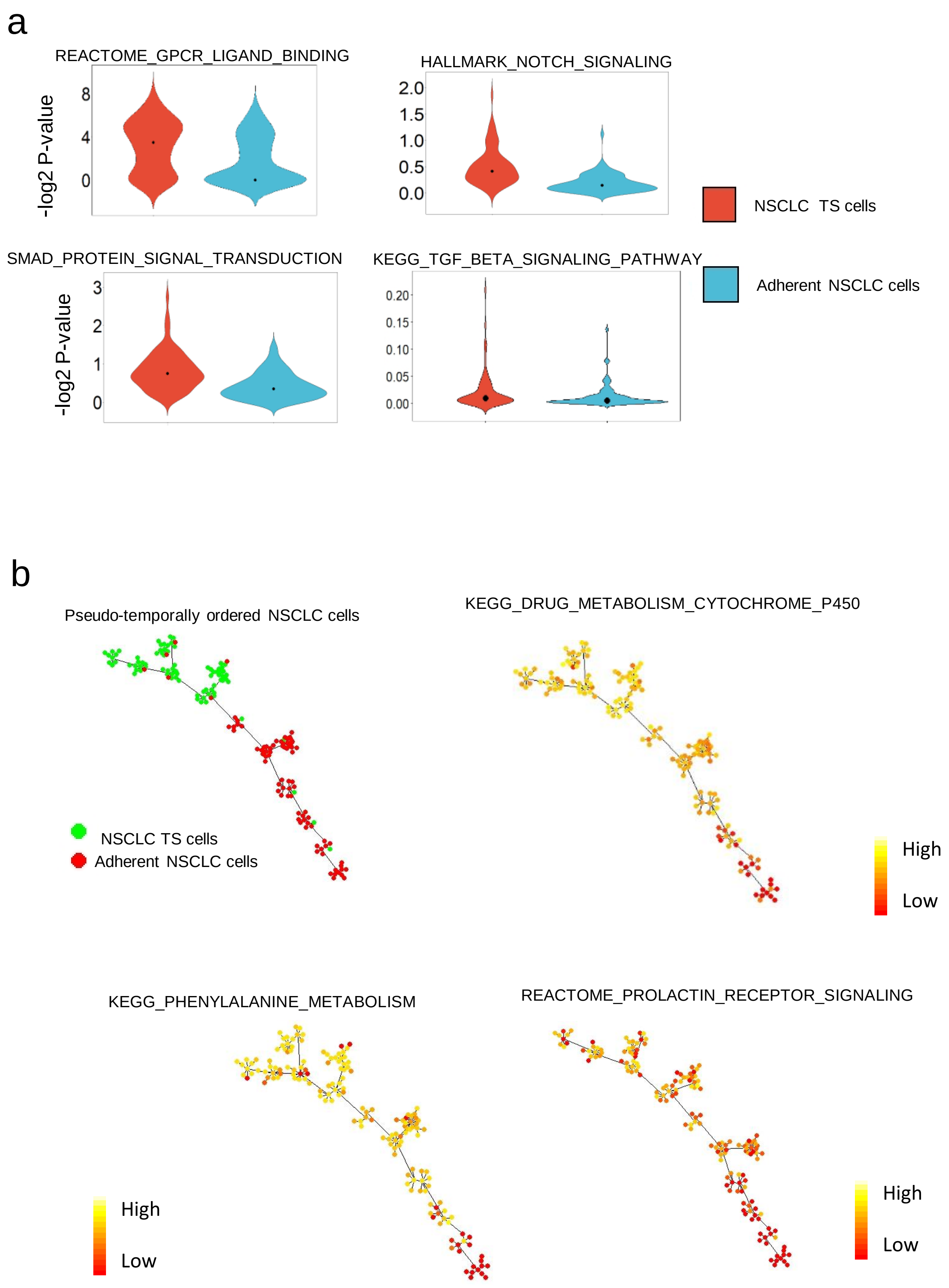


**Figure S13**: Visualization of distribution and gradient of enrichment score of pathways in non-small cell lung cancer (NSCLC) cells (a) Violin plot of enrichment scores for 4 pathways in Adh and TS cells of NSCLC. NOTCH, GPCR and SMAD signalling showed a significant difference (P-value < 0.01) in enrichment scores in TS and Adh cells. Whereas TGF-beta signalling pathway scores remained similar in both cell lines. (b) Pseudo-temporal ordering of lung cancer cells. The gradient of pathway scores for three different pathways/gene-sets are also shown.
